# Plant YTHDF proteins are direct effectors of antiviral immunity against an m^6^A-containing RNA virus

**DOI:** 10.1101/2022.10.19.512835

**Authors:** Mireya Martínez-Pérez, Frederic Aparicio, Laura Arribas-Hernández, Mathias Due Tankmar, Sarah Rennie, Peter Brodersen, Vicente Pallas

## Abstract

In virus-host interactions, nucleic acid-directed first lines of defense that allow viral clearance without compromising growth are of paramount importance. Plants use the RNA interference pathway as such a basal antiviral immune system, but additional RNA-based mechanisms of defense also exist. The infectivity of the plant positive strand RNA virus alfalfa mosaic virus (AMV) relies on demethylation of viral RNA by recruitment of the cellular *N6*-methyladenosine (m^6^A) demethylase ALKBH9B, but how demethylation of viral RNA promotes AMV replication remains unknown. Here, we show that inactivation of the cytoplasmic YT521-B homology domain (YTH)-containing m^6^A-binding proteins, ECT2, ECT3, and ECT5 is sufficient to restore AMV infectivity in partially resistant *alkbh9b* mutants. We also show that the antiviral function of ECT2 is distinct from its previously demonstrated function in promotion of proliferation of primordial cells, because an ECT2 mutant carrying a small deletion in its intrinsically disordered region is partially compromised for antiviral defense, but not for developmental functions. These results indicate that the m^6^A-YTH axis constitutes a novel branch of basal antiviral immunity in plants.

## INTRODUCTION

Viruses are intracellular pathogens that use their genetic code to turn cells into their multiplication factories. To defend against hostile take-over by viral genetic material, host organisms have evolved mechanisms to recognize foreign nucleic acids. For example, bacteria use a combination of restriction-modification and CRISPR-Cas systems to eliminate viral nucleic acids (Barrangou *et al*, 2007; Loenen *et al*, 2014; Egido *et al*, 2022), and a combination of cyclic nucleotide signaling and NAD+ depletion systems coupled to nucleic acid sensors to eliminate infected cells from a population (Tal *et al*, 2021; Koopal *et al*, 2022). Similar basal antiviral immune systems operating directly on viral nucleic acids have been found in animals where at least four distinct cytoplasmic systems recognize double-stranded features of viral RNA to activate signaling pathways that lead to induction of innate antiviral immune responses (Rehwinkel & Gack, 2020; Wu & Chen, 2014; Holleufer *et al*, 2021; Slavik *et al*, 2021). In plants, the double-strandedness of cytoplasmic viral RNA is also used to distinguish it from endogenous RNA, since plants use RNA interference (RNAi) as a potent basal antiviral defense mechanism (Lindbo, 2012). In addition, extracellular dsRNAs can be sensed by as yet unidentified receptors to effect antiviral pattern-triggered immunity PTI (Niehl & Heinlein, 2019; Meier *et al*, 2019).The effectiveness of RNAi as a basal antiviral defense mechanism is illustrated by the fact that nearly all plant viruses use anti-RNAi effectors as virulence factors (Jin *et al*, 2021; Csorba *et al*, 2015; Voinnet *et al*, 1999). In plants, RNAi remains the most generally employed basal antiviral defense mechanism to date, but features of viral genetic material other than its double-strandedness may be used for distinction between cellular and viral RNA, and hence, for antiviral defense. For example, in both mammals and plants, non-sense-mediated decay may detect and repress long viral RNAs with stop codons of reading frames located towards the 5’-end of the viral transcript (Garcia *et al*, 2014; Balistreri *et al*, 2014); a feature that viruses may circumvent either through the use of anti-NMD factors or by organizing their genomes into a long poly-protein-encoding mRNA containing only a single downstream stop codon (Popp *et al*, 2020; Balistreri *et al*, 2014). Furthermore, in the nematode *C. elegans* that also relies on RNAi for basal antiviral defense (Gammon, 2017), a terminal uridyltransferase exerts antiviral immunity independently of RNAi, although the feature of viral RNA recognized in this case remains unclear (Le Pen *et al*, 2018).

Covalent modification of mRNA is now understood to play important roles in regulation of gene expression, and many viruses contain modified nucleotides of importance for the outcome of the host-virus interaction (Courtney, 2021; Gokhale & Horner, 2017). The most widespread, and probably most important, modification of eukaryotic cellular mRNA is *N*^*6*^-methyladenosine (m^6^A) (Wiener & Schwartz, 2021; Arribas-Hernández & Brodersen, 2020). m^6^A is installed at DRACH (D=A/G/U, R=G/A, H=A/C/U) or GGAU motifs in pre-mRNA by a dedicated, nuclear multi-subunit methyltransferase, or “writer”, complex (Arribas-Hernández & Brodersen, 2020; Arribas-Hernández *et al*, 2021a). This highly conserved complex consists of a heterodimeric catalytic core of methyltransferase-like proteins known as MAC, and a large MAC-associated complex (MACOM) required for activity *in vivo*. In Arabidopsis, MAC consists of MTA (AT4G10760, orthologous to metazoan Mettl3) and MTB (AT4G09980, orthologous to metazoan Mettl14), while MACOM is composed of several factors, including FKBP12 INTERACTING PROTEIN 37 (FIP37, AT3G54170, orthologous to metazoan WTAP), VIRILIZER (VIR, AT3G05680), the ubiquitin ligase HAKAI (AT5G01160), and the Zn-finger protein HAKAI-INTERACTING Zn-FINGER2 (HIZ2; AT5G53440, equivalent to metazoan ZC3H13/Flacc) (Zhong *et al*, 2008; Shen *et al*, 2016; Růžička *et al*, 2017; Zhang *et al*, 2022). m^6^A can be demethylated, or “erased”, by a set of enzymes belonging to the AlkB family of Fe(II)- and α-ketoglutarate-dependent dioxygenases (Van den Born *et al*, 2009). In Arabidopsis, the AlkB family is composed of 14 proteins (Mielecki *et al*, 2012; Kawai *et al*, 2014). Demethylation activity has been demonstrated experimentally for ALKBH9B and ALKBH10B (Mielecki *et al*, 2012; Kawai *et al*, 2014; Duan *et al*, 2017; Martínez-Pérez *et al*, 2017), and recent evidence suggests its presence also in ALKBH9C (Amara *et al*, 2022).

It is an important function of m^6^A to generate a binding site for RNA binding proteins specialized for the m^6^A recognition via a so-called YT521-B homology (YTH) domain (Imai *et al*, 1998; Zhang *et al*, 2010; Liao *et al*, 2018). The YTH-domain contains an aromatic cage that forms a hydrophobic pocket for the methyl group in m^6^A, thus resulting in 10-20-fold higher affinity for m^6^A-containing RNA than for unmethylated RNA (Theler *et al*, 2014; Patil *et al*, 2018; Xu *et al*, 2014, 2015; Zhu *et al*, 2014; Li *et al*, 2014). For that reason, YTH domain proteins are sometimes referred to as m^6^A readers. In the cytoplasm, a family of so-called YTHDF proteins binds to m^6^A-containing RNA, and while genetics shows these YTHDF proteins to be of key importance for plant and animal development (Arribas-Hernández *et al*, 2020, 2018; Lasman *et al*, 2020; Kontur *et al*, 2020), the exact reasons for this biological importance remain undefined in plants, and heavily debated in mammals (Murakami & R.Jaffrey, 2022; Tsutsui & Higashiyama, 2017; Lasman *et al*, 2020). In the flowering plant Arabidopsis, there are 11 YTHDF proteins, named EVOLUTIONARILY CONSERVED C-TERMINAL REGION (ECT) because of the presence of the conserved YTH domain in the C-terminal part, after an N-terminal intrinsically disordered region (IDR) as in YTHDF proteins of other organisms. Only ECT2, ECT3, and ECT4 have been shown to work as m^6^A readers (Arribas-Hernández *et al*, 2018; Scutenaire *et al*, 2018; Wei *et al*, 2018). In Arabidopsis, the YTHDF proteins ECT2 and ECT3 act redundantly to stimulate cellular proliferation in organ primordia, such that *ect2/ect3* double mutants show delayed leaf and flower formation, slow root and stem growth, and aberrant leaf, flower, silique, and trichome morphology (Arribas-Hernández *et al*, 2020, 2018). In nearly all cases, these phenotypes are exacerbated by additional mutation of *ECT4* (Arribas-Hernández *et al*, 2020, 2018). Recent studies report similar roles of rice and tomato YTHDF proteins in plant development (Yin *et al*, 2022; Ma *et al*, 2022).

m^6^A is also found in viral mRNAs, first seen as early as 1975 in simian virus 40 mRNAs (Lavi & Shatkin, 1975). Subsequent studies have shown presence of m^6^A in viral RNAs from several mammalian RNA and DNA viruses (Baquero-Perez *et al*, 2021; Wu *et al*, 2020; Williams *et al*, 2019), and direct binding of YTH proteins to RNAs from some of these viruses, such as hepatitis C virus (HCV), zika virus (ZIKV) or human immunodeficiency virus-1 (HIV-1) has been demonstrated (Gokhale *et al*, 2016; Kennedy *et al*, 2016; Lichinchi *et al*, 2016). In plant viruses, fewer examples of the involvement of m^6^A in infection cycles have been described (Martínez-Pérez *et al*, 2017, 2021; van den Born *et al*, 2008; Tian *et al*, 2021; Zhang *et al*, 2021).

We previously showed that viral RNA from a positive strand plant RNA virus, alfalfa mosaic virus (AMV), contains m^6^A, and that AMV infectivity of wild type Arabidopsis plants depends on recruitment of a cellular m^6^A demethylase, ALKBH9B, to viral RNA (Martínez-Pérez *et al*, 2017). In *alkbh9b* knockout plants, local and systemic AMV infections are attenuated and, most remarkably, viral invasion of the floral stems is almost blocked. Thus, methylation of the viral RNA is part of an important defense mechanism that the infectious virus adapts to by manipulation of ALKBH9B to act on the viral RNA (Alvarado-Marchena *et al*, 2021; Martínez-Pérez *et al*, 2021). The molecular basis of m^6^A-dependent antiviral defense remains undefined, however. It may, for example, involve m^6^A-binding reader proteins or RNA structure properties related to the weaker m^6^A-U than A-U base pairs, the latter perhaps suggested by the observation that adenosines in a DRACH context within the 3’UTR of alfalfa mosaic virus (AMV) RNA 3 represent a key structural requirement for viral replication (Alvarado-Marchena *et al*, 2022).

Here, we show that mutation of the *ECT2/ECT3/ECT4/ECT5* module in Arabidopsis reduces AMV resistance, and that the increased AMV resistance of Arabidopsis *alkh9b* mutants can be reverted by mutation of *ECT2/ECT3/ECT5*. We also show that the functions of ECT2 in stimulation of cellular proliferation and antiviral defense can be separated by a small deletion in its N-terminal intrinsically disordered region (IDR). These results establish the m^6^A-YTHDF axis as a novel basal antiviral defense layer.

## RESULTS

### AMV infection induces components of the m^6^A-YTH axis

To guide genetic dissection of m^6^A-mediated defense against AMV, we first analyzed transcriptome changes upon AMV infection with particular attention to transcripts encoding components of the m^6^A pathway. For this experiment, we selected young, emerging rosette leaves of infected plants, since m^6^A binding proteins and methyltransferase subunits were shown to be mostly expressed in tissues with high cell division rates (Arribas-Hernández *et al*, 2020, 2018; Zhong *et al*, 2008). Differential expression analysis showed that 2611 genes were significantly upregulated (log_2_FC≥1; FC, fold change), whereas only 194 genes were downregulated (log_2_FC≤-1) during AMV infection (**Figure 1A**). Molecular functions among differentially expressed genes were enriched in protein-protein interaction and DNA- or RNA-binding activities (**Figure 1B**). This set also included some components of the m^6^A machinery (**Figure 1C**). The levels of mRNAs encoding three components of the methylation complex – *MTA, MTB*, and *VIR* – were upregulated by the infection, whereas the levels of mRNAs encoding potential m^6^A erasers did not substantially change (**Figure 1C**). Among YTHDF-encoding genes, only *ECT5* showed a greater than 2-fold upregulation (**Figure 1C**). We verified the induction of *ECT5*, along with that of *MTA, MTB*, and *VIR*, using quantitative RT-PCR analysis (**Figure 1D**). The previously described functional m^6^A readers ECT2 and ECT3 (Arribas-Hernández *et al*, 2018) were also significantly upregulated, albeit with smaller effect size (log_2_FC values 0.6 and 0.4 for *ECT2* and *ECT3*, respectively; **Figure 1C**).

**Figure 1.**
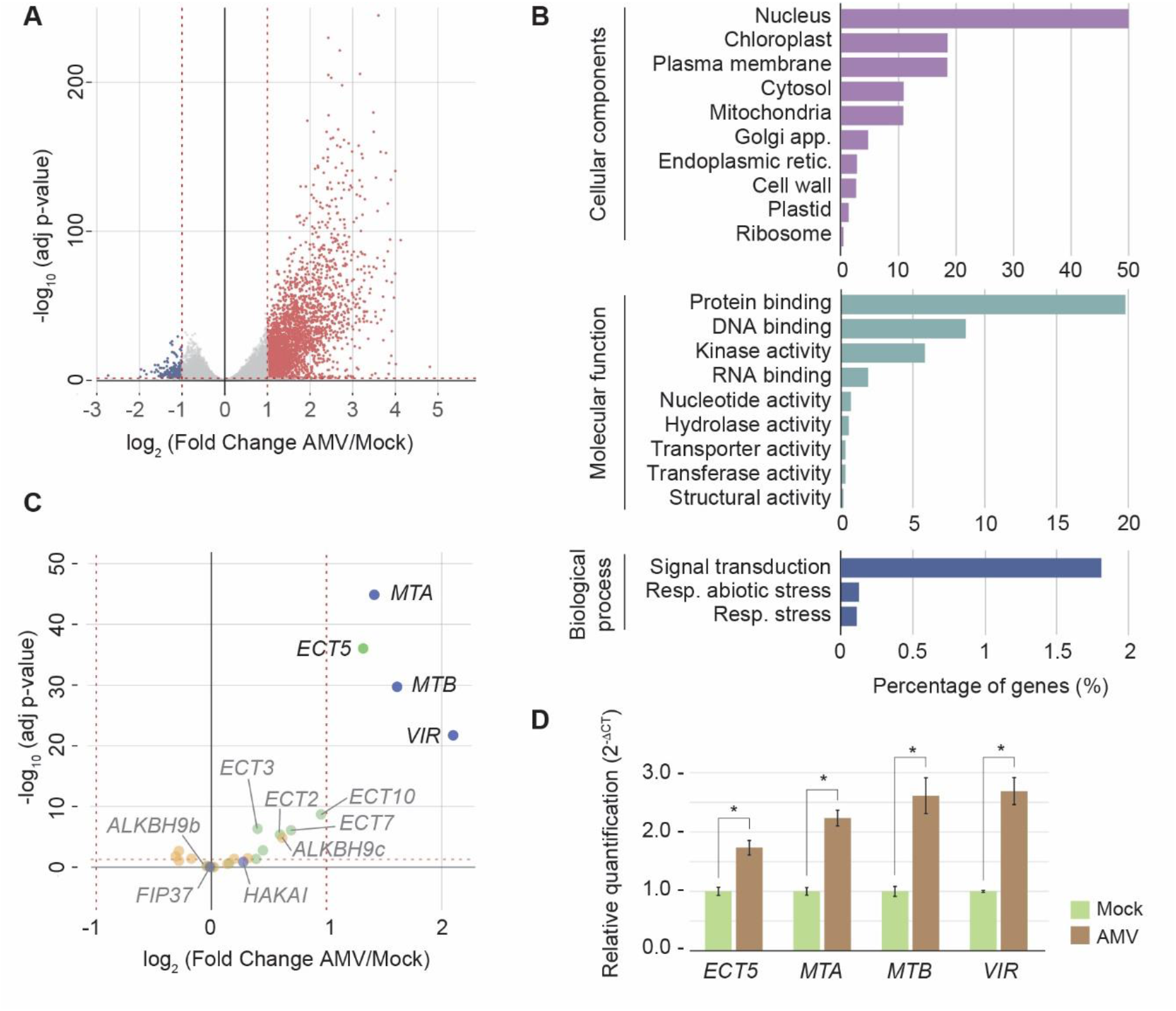
Differential gene expression analysis between mock and AMV-infected Arabidopsis plants. **(A)** Volcano plot depicting differential expression of nearly 18000 unique genes (minimum expression of 1 read per kilobase per million, RPKM) in response to AMV infection. In red, upregulated genes (log_2_FC ≥ 1; adjusted p-value ≤ 0.05); in blue, downregulated genes (log_2_FC ≤ -1; adjusted p-value ≤ 0.05). **(B)** Most highly enriched gene ontology (GO) terms in the set of differentially expressed genes. **(C)** Filtered volcano plot visualizing the expression of genes involved in the m^6^A pathway in response to AMV infection. **(D)** RT-qPCR analysis of expression of four upregulated m^6^A pathway-related genes in response to AMV infection (three biological replicates). Error bars represent the standard error of the mean (SEM). Asterisks indicate a *p* < 0.05 applying Student’s t-test for ΔCt mean values (n = 3).

We next analyzed single *ect2-1, ect3-1, ect4-2* mutants (Arribas-Hernández *et al*, 2018), and an *ect5* T-DNA insertion mutant (*ect5-1*, SALK_131549; **Supplemental Figure 1A, B**) for possible defects in AMV resistance (**Supplemental Table 1**). No single mutants showed significant differences in AMV titers (**Supplemental Figure 1C, D**), perhaps due to functional redundancy as demonstrated for their growth-promoting functions (Arribas-Hernández *et al*, 2020). We therefore analyzed composite mutants of *ECT2, ECT3*, and *ECT4*. The results showed clearly elevated AMV titers in non-inoculated leaves (systemic infection), but not in local infection contexts, of AMV-infected *ect2-1/ect3-1* (henceforth, *de23*) and *ect2-1/ect3-1/ect4-2* (henceforth, *te234*) mutant plants compared to wild type (**Figure 2A-B**). Double mutants involving *ect4* showed weak, if any, effects on AMV accumulation, while *te234* tended to have somewhat higher AMV titers than *de23*, although this effect was variable in size between experiments (**Figure 2B**; **Supplemental Figure 2B**). The differences in systemic viral load between wild type and *ect2/ect3* double knockouts were confirmed using an *ect2/ect3* double mutant with independent knockout alleles (*ect2-3/ect3-2*, **Supplemental Figure 2**; **Supplemental Table 1**) (Arribas-Hernández *et al*, 2018). Next, since *ECT5* is tightly linked to *ECT2*, we used CRISPR-Cas9 to produce the double *ect2-1/ect5-2* (*de25*) and triple *ect2-1/ect3-1/ect5-4* (*te235*) mutants (**Supplemental Figure 3**). Importantly, we observed increased AMV titers in non-inoculated, but not in locally infected, tissues of *de25* compared to WT (**Figure 2C-D**), pointing to an important involvement of ECT5 in limiting systemic AMV infection. In contrast to *de23* mutants, *de25* mutants exhibited no obvious leaf morphology defects or delay in leaf formation (**Figure 2E**). As observed for the *te234* mutants, *te235* tended to show moderately higher AMV titers systemically than *de23*, although with some variability between experiments (**Figure 2D, Supplemental Figure 2C**). Similar results were obtained with different CRISPR-induced *ect5* alleles (**Supplemental Figure 4; Supplemental Table 1**). We conclude that ECT2, ECT3, and ECT5 are required for systemic resistance to AMV, observable in *ect2/ect3* and *ect2/ect5* mutants, and that ECT4 seems to have a minor role as its effect is mostly observable as a variable enhancement of the increased susceptibility of *ect2/ect3* double knockout mutants in non-inoculated aerial tissues.

**Figure 2.**
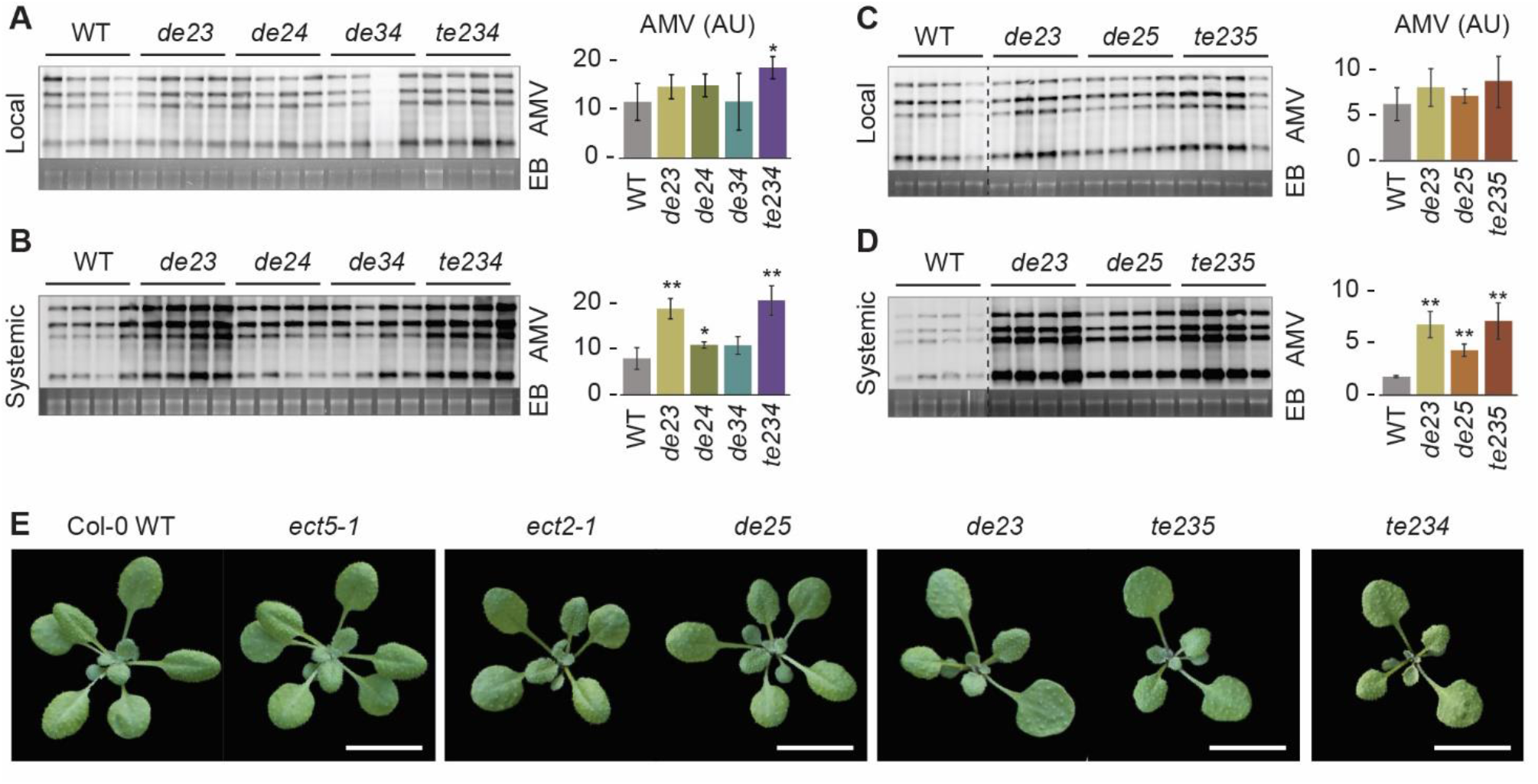
Combined inactivation of *ECT2/3/4/5* enhances titers of AMV in systemic tissues. **(A-D)** RNA blots were hybridized to AMV-specific probes to reveal accumulation of viral RNA in double (de) and triple (te) *ect2/3/4* (A, B) or *ect2/3/5* (C, D) mutants compared to WT plants. A and C show local infection at 3 dpi; B and D show systemic infection in aerial tissue at 7 dpi. Each panel shows a representative RNA blot displaying AMV RNAs 1-4 (left) and its quantification histogram (right). Dashed lines indicate non-contiguous samples analyzed on the same membrane. Ethidium bromide staining of ribosomal RNAs (EB) was used as RNA loading control. Error bars indicate standard deviations. Asterisks indicate significant differences from the WT (*: *p* < 0.05; **; *p* < 0.01) using the Student’s t-test (n = 4). AU, arbitrary units. **(E)** Phenotypes of rosettes of the indicated genotypes at 19 days after germination. Scale bars: 1 cm.

### The m^6^A-binding activity of ECT2 is required for its antiviral activity

The amino acid residues of human YTHDF proteins implicated in RNA and m^6^A binding are highly conserved in Arabidopsis ECT1-4 proteins. This includes in particular the three tryptophan residues that form the m^6^A-binding aromatic cage (Arribas-Hernández *et al*, 2018). Mutational studies of these tryptophan residues have demonstrated that integrity of the aromatic cage is required for ECT2 binding to m^6^A-modified RNAs (Scutenaire *et al*, 2018) and for ECT2, ECT3, and ECT4 function *in vivo* (Arribas-Hernández *et al*, 2018, 2020). Thus, to test whether ECT-mediated antiviral resistance requires the m^6^A-binding activity, we used *de23* mutants transformed with either *ECT2*^*WT*^*-mCherry* or with the m^6^A-binding deficient point mutant *ECT2*^*W464A*^*-mCherry* (**Supplemental Table 1**). The ECT2^W464A^-mCherry-expressing lines were clearly defective in AMV resistance, while expression of ECT2^WT^-mCherry in *de23* restored resistance to levels comparable to that of wild type or *ect3-1* (**Figure 3A**, upper and lower panels show two independent complementation transgenic lines).

**Figure 3.**
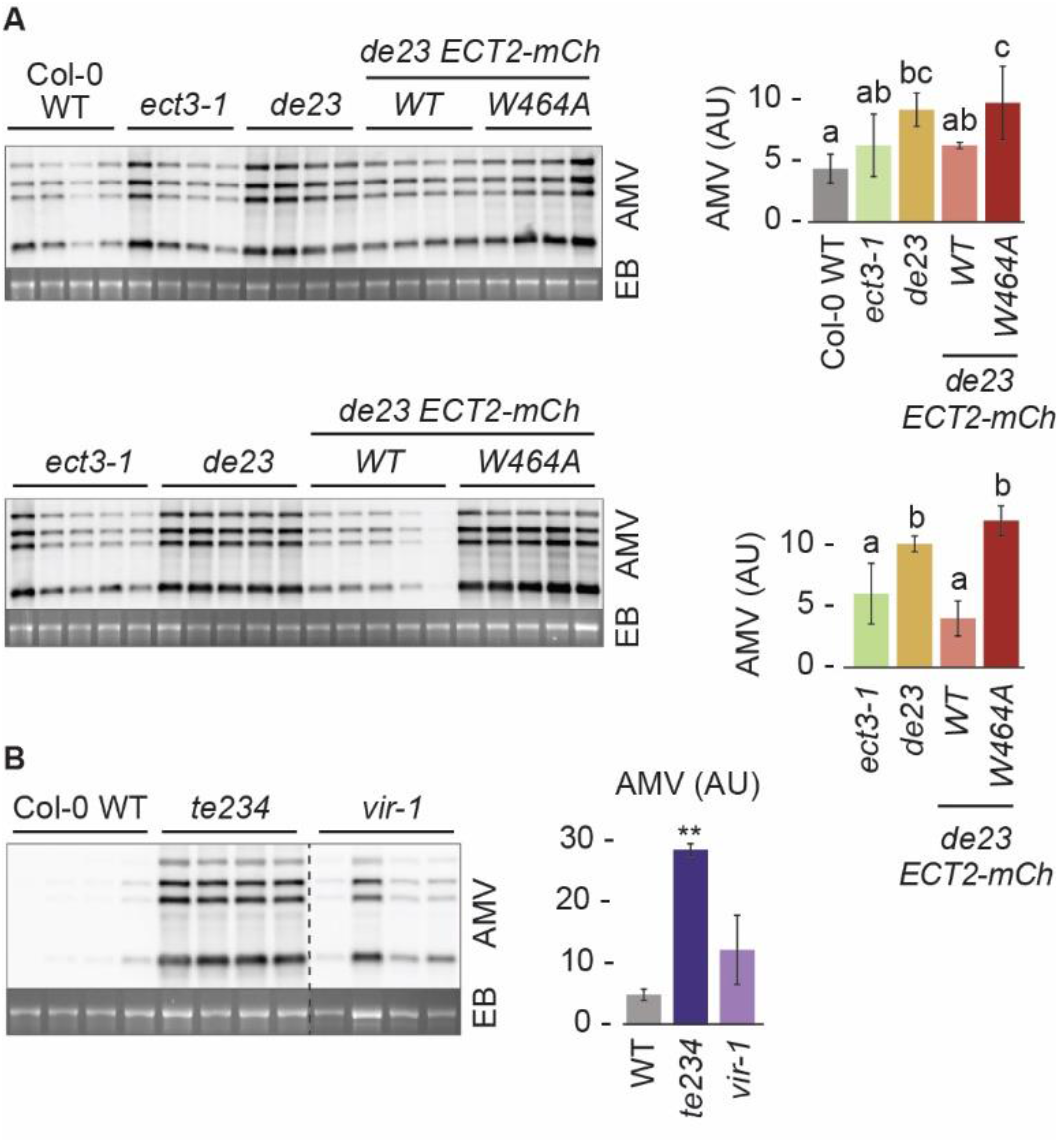
ECT-mediated defense against AMV requires intact m^6^A-binding pockets and is not an indirect effect of a dysfunctional m^6^A-ECT axis. **(A-B)** RNA-blot analysis of AMV systemic infection in plants expressing ECT2^WT^-mCherry (WT) or ECT2^W464A^-mCherry (W464A) in the *de23* background (A) and in *vir-1* plants (B) at 7 dpi. Each panel shows a representative RNA blot displaying AMV RNAs 1-4 (left) and its quantification histogram (right). Ethidium bromide staining of rRNAs (EB) was used as RNA loading control. Genotypes are indicated on the top of each northern blot. In A, upper and lower panels show independent complementation transgenic lines, and different letters indicate statistical differences according to Fisher’s Least Significant Difference (LSD) (*p* < 0.05). In B, dashed line indicates non-contiguous samples that are analyzed on the same membrane. Error bars indicate standard deviations. Asterisks indicate significant differences from the WT (*: *p* < 0.05; **: *p* < 0.01) applying Student’s t-test (n = 4). AU, arbitrary units.

We note that the loss of AMV resistance in both knockout and ECT2^W464A^-mCherry-expressing lines could in principle be an indirect consequence of the developmental defect of *ect* mutants that may render the plant more amenable to viral replication rather than a direct effect of missing or defective association with m^6^A-containing viral RNA. We disfavor this possibility for two reasons. First, in contrast to *de23, de25* mutants display no obvious rosette phenotype (**Figure 2E**), yet the increase in AMV titers compared to wild type is robust (**Figure 2D**). Second, mutation of the nuclear m^6^A writer component *VIR* causes a rosette phenotype similar to, but slightly stronger than *ect2/ect3/ect4* (Růžička *et al*, 2017), yet does not lead to the same degree of loss of antiviral resistance (**Figure 3B**).

### ECT2 associates with AMV RNA *in vivo*

We next sought to establish whether ECTs associate with AMV RNA, focusing specifically on ECT2 due to its higher expression levels compared to ECT3 and ECT5 (Arribas-Hernández *et al*, 2021b). We used the proximity-labeling method HyperTRIBE (Targets of RNA-binding proteins Identified by Editing) (McMahon *et al*, 2016), in which the catalytic domain of the fly Adenosine Deaminase Acting on RNA (ADAR) is fused to ECT2 (**Supplemental Table 1**) and expressed under the control of ECT2 promoter. Because adenosine-to-inosine editing by ADAR is detected as A-to-G transitions by RNA-Seq, RNA bound by ECT2 can be identified by virtue of its higher A-G editing proportions compared to control lines expressing unfused ADAR (McMahon *et al*, 2016; Xu *et al*, 2018; Rennie *et al*, 2021). Using an experimental set-up similar to our previous HyperTRIBE analysis which faithfully identified direct endogenous mRNA targets of ECT2 and ECT3 (Arribas-Hernández *et al*, 2021a, 2021b), we employed five independent transgenic lines expressing either the ECT2-FLAG-ADAR fusion or the free FLAG-ADAR, the latter acting as negative controls. The negative control lines showed on average higher FLAG-ADAR expression levels than the ECT2-FLAG-ADAR fusion lines (**Figure 4A**), contributing to higher stringency in the analysis, although in contrast, viral titers were generally lower in these same lines, thus providing fewer sequenced reads for the viral RNAs (**Figure 4B)**. Furthermore, nearly all detected AMV sequence reads mapped to the positive strand, as expected (**Figure 4C**). We first selected 395 candidate positions showing A-G changes compared to the reference AMV sequence in at least two of five lines. However, since the viral RNA-dependent RNA polymerase introduces random mutations, and since free ADAR has a non-zero background editing activity, we sought to formally test whether significantly higher editing proportions were observed in ECT2-FLAG-ADAR lines compared to FLAG-ADAR control lines. For this purpose, we employed our recently published method for the analysis of HyperTRIBE data (Rennie et al., 2021) identifying two sites in the positive strand of RNA2 (RNA2_701 and RNA2_2271) that show consistent A-G changes across FLAD-ADAR fusion lines, with modeled editing proportions significantly higher than in FLAG-ADAR control lines (FDR < 0.05, **Figure 4D**). We further noted an additional likely *bona fide* site on RNA1 (RNA1_2482), which did not make it under the FDR threshold, likely due to lack of power resulting from low editing proportions. In all three cases, editing proportions in the ECT2-FLAG-ADAR fusion lines strongly correlated with corresponding ECT2-FLAG-ADAR expression levels across the five individual transgenic lines used as replicates (**Figure 4E**; **Supplemental Figure 5**), supporting the conclusion that these A-G changes are a result of editing by the ECT2-FLAG-ADAR fusion. Finally, we noted that while the fused ADAR to ECT2 is not expected to act on the exact binding site, the two high-confidence sites in RNA2 did appear in the vicinity of DRACH motifs, which may be indicative of the methylated position to which ECT2 was recruited (**Figure 4F, G**). Taken together, our HyperTRIBE data indicate that ECT2 associates with AMV RNA *in vivo*.

**Figure 4.**
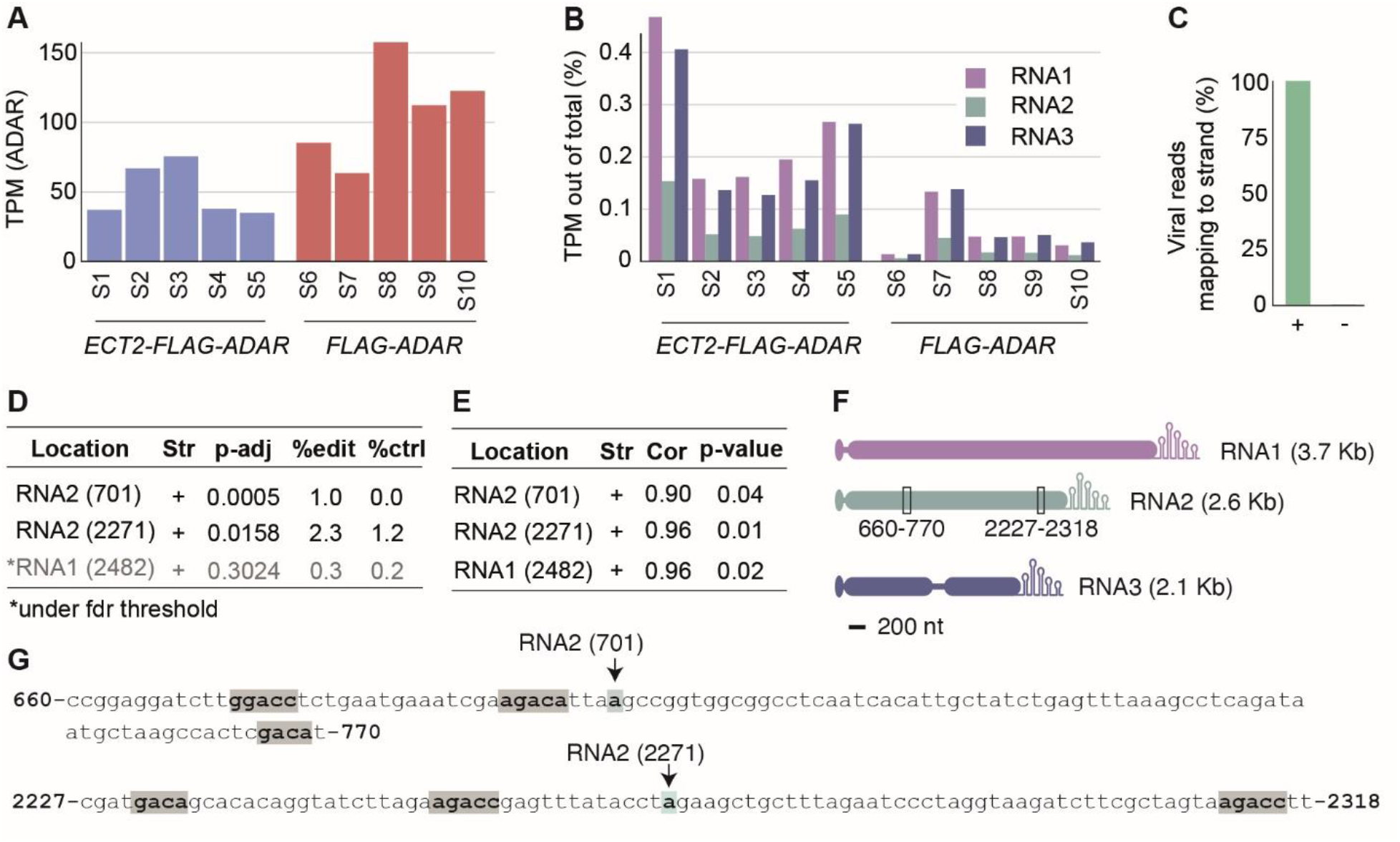
HyperTRIBE identifies *in vivo* interactions between ECT2 and AMV RNAs. **(A)** ADAR expression (TPM – Transcripts Per Kilobase Million – mapping) in five samples of ECT2-FLAG-ADAR (S1-S5) and FLAG-ADAR (S6-S10) transgenic lines. **(B)** Percentage of total reads mapping to the AMV RNAs in the different samples. **(C)** Percentage of reads mapping to either the positive or negative AMV strand. **(D)** Positions preferentially edited in AMV RNAs of the ECT2-FLAG-ADAR compared to FLAG-ADAR lines. %edit and %ctrl show the editing proportion (G/(A+G)) in the ECT2-FLAG-ADAR and FLAG-ADAR lines, respectively. Significant sites were determined using hyperTRIBE**R** pipeline (Rennie *et al*, 2021). To be formally tested, an A-to-G change had to be present in at least two of the five ECT2-FLAG-ADAR samples. **(E)** Correlation between ADAR log TPM and editing proportions across individual ECT2-FLAG-ADAR transgenic lines for the three most highly significantly edited sites. **(F)** Schematic representation of the positions of the two high-confident ECT2-dependent editing sites in AMV RNAs. The editing sites nearby DRACH motives are highlighted in light blue and gray, respectively.

### The m^6^A-ECT axis constitutes a basal layer of antiviral defense

The observations that inactivation of *ECT2/3/(4)/5* leads to higher AMV infectivity, and that ECT2 directly associates with AMV RNA *in vivo*, are consistent with a model in which binding of ECTs to viral RNA causes inhibition of viral replication. However, the results reported thus far suffer from the limitation that they do not analyze the true effect of ECT association with AMV RNA, because they are carried out with a wild type virus in an otherwise wild type *Arabidopsis* genetic background. Since AMV requires the endogenous demethylase ALKBH9B, recruited probably via its capsid protein (CP) to AMV RNA, for infectivity (Martínez-Pérez *et al*, 2017), these experiments address the influence of ECTs on infection in a setting in which a virulence mechanism is diminishing the impact of adenosine methylation of viral RNA. Because the CP has multiple essential functions in the infection cycle, including RNA encapsidation and initial translational control (Herranz *et al*, 2012), informative genetic manipulation of the CP is not straightforward. By contrast, our previous study showed that the systemic infection in the floral stems is nearly fully blocked upon disruption of the host *ALKBH9B* gene. We therefore constructed *alkbh9b/te235* mutants (**Supplemental Table 1**) to test the impact of *ECT* inactivation in a setting in which AMV infectivity is strongly incapacitated due to hypermethylation. The AMV titers in *alkbh9b/te235* were intermediate between *alkbh9b* and *te235* at early stages of infection (7 days post-inoculation, dpi) (**Figure 5A**), but were identical to *te235* (and Col-0) at late stages (20 dpi) (**Figure 5B**), where little to no AMV was detectable in *alkbh9b*. We interpret these results to mean that m^6^A-ECTs efficiently limit the speed of systemic spread of AMV to the point where nearly no virus accumulates in stems upon leaf inoculation in *alkbh9b* (Figure 5B) (Martínez-Pérez *et al*, 2017, 2021), even at late stages of infection. At such late points, additional aspects of the host-AMV interaction are likely to limit AMV titers to a maximal value reached by both Col-0 and *te235*. Although spreading and systemic accumulation is lower in *alkbh9b/te235* than in *te235*, the virus eventually reaches these same maximal titers systemically, in marked contrast to what is observed in *alkbh9b*. Importantly, therefore, inactivation of *ECT2/3/5* is sufficient to break the strong m^6^A-based protection against systemic infection in the absence of ALKBH9B, clearly showing that ECT2/3/5 are requisite effectors of the m^6^A-based defense against AMV. These results argue that for AMV, and perhaps other viruses whose RNA is *N6*-adenosine methylated, the m^6^A-ECT axis constitutes a first, basal layer of antiviral defense that must be overcome for the virus to be infectious. Indeed, the results conceptually parallel the restoration of infectivity of RNA virus strains with defective anti-RNAi effectors by inactivation of the RNAi machinery (Deleris *et al*, 2006; Wang *et al*, 2011).

**Figure 5.**
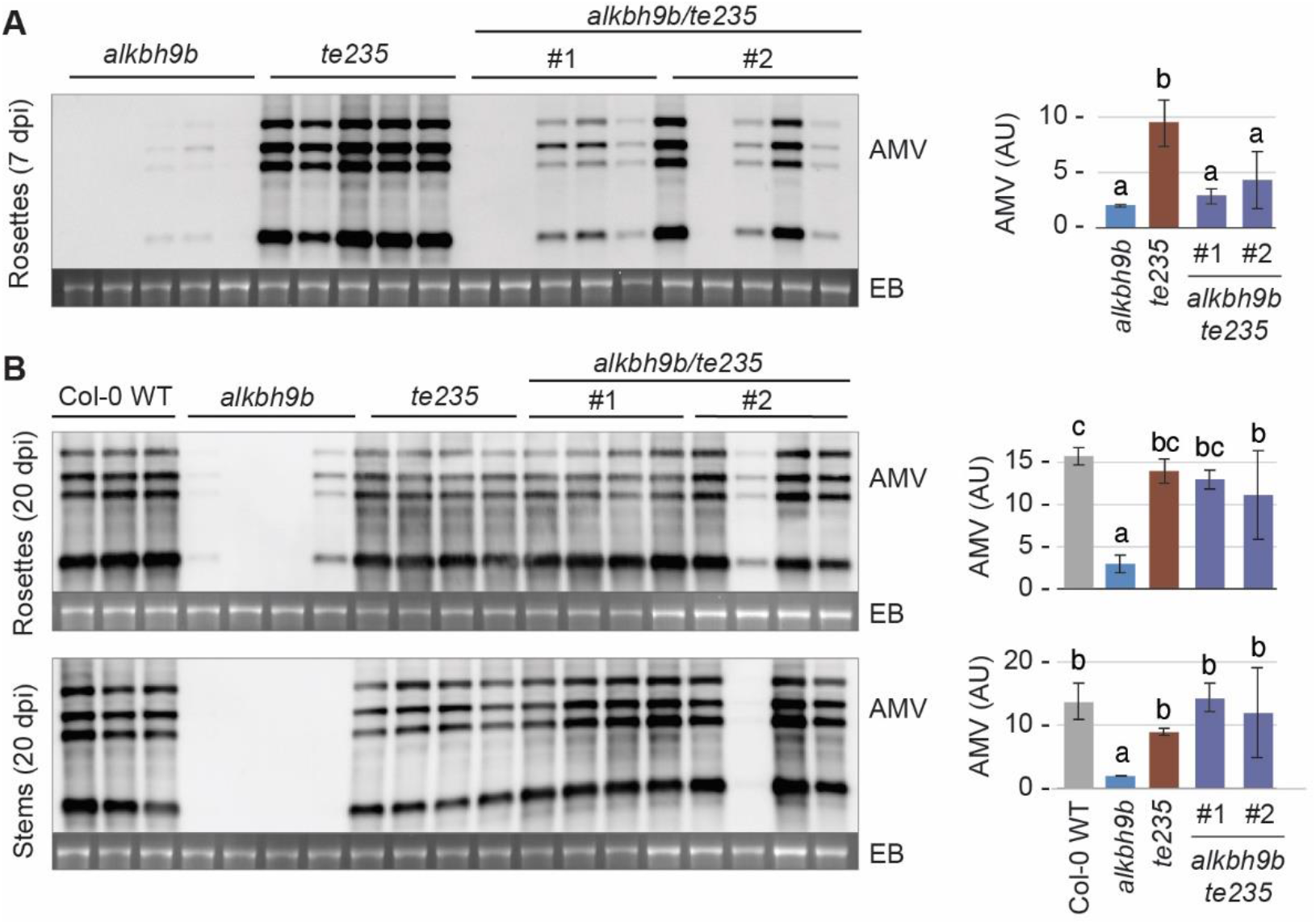
The ECT2/3/5 module constitutes a basal defense layer against AMV infection, since knockout of *ECT2/3/5* in the *alkbh9b* background recovers susceptibility. **(A-B)** Systemic infection of rosettes at 7 dpi (A) or rosettes and floral stems at 20 dpi (B). Each panel shows a representative RNA blot displaying AMV RNAs 1-4 (left) and its quantification histogram (right). Ethidium bromide staining of rRNAs (EB) was used as RNA loading control. Genotypes are indicated on the top of each northern blot. #1 and #2 correspond to the progeny of two different quadruple homozygous F2 siblings derived from the *alkbh9b* x *te235* cross. Error bars show standard deviations. AU, arbitrary units. Different letters indicate statistical differences according to Fisher’s Least Significant Difference (LSD) (*p* < 0.05).

### The growth-promoting and antiviral activities of ECT2 are genetically separable

The model of basal antiviral defense mediated by m^6^A-ECT2/3/4/5 may be at odds with the described endogenous function of these factors in promoting cellular proliferation (Arribas-Hernández *et al*, 2020), as a highly metabolically active state favors viral replication. Thus, it is difficult to imagine how the very same biochemical activity would promote both antiviral defense and cellular proliferation. We therefore reasoned that ECTs may use different biochemical and/or biophysical properties to achieve both effects. To analyze this possibility, we made use of a series of small deletions engineered in the long N-terminal intrinsically disordered region (IDR). These deletions were originally designed to identify regions of importance for the developmental function of ECT2 and will be described in detail in a later communication. Of six ∼50-80 aa deletions, four had no effect on leaf formation. Since the presence of Tyr residues has been described as a crucial sequence feature for liquid-liquid phase separation in different proteins (Wang *et al*, 2018; Martin *et al*, 2020; Bremer *et al*, 2022), we selected the deletion ECT2^ΔN5^ featuring a Tyr-rich region present in many ECTs for analysis of AMV resistance (**Figure 6A; Supplemental Figure 6A, B**). We observed a reduction in AMV resistance compared to ECT2^WT^ when both were expressed in the *te234* mutant background (**Figure 6B**). This effect was reproducible, albeit somewhat variable, across several transgenic lines that all expressed similar levels of ECT2-mCherry or ECT2^ΔN5^-mCherry (**Figure 6C**). Because the rate of leaf formation and leaf morphology in *te234* lines expressing ECT2^ΔN5^-mCherry were indistinguishable from those expressing ECT2^WT^-mCherry (**Figure 6D, E; Supplemental Figure 6C**), yet AMV resistance was measurably compromised, these results suggest that the IDR of ECT2 harbors two separable activities employed to achieve different goals: one that stimulates cellular proliferation by binding to endogenous m^6^A-containing mRNA, and one that effects basal antiviral resistance when ECT2 binds to hypermethylated viral RNA. We note that since the antiviral activity was not fully abolished in *te234* expressing ECT2^ΔN5^-mCherry, it is formally possible that the different impacts of this mutation on antiviral defense and developmental phenotypes are caused by a threshold effect, such that the same reduction in molecular activity would give rise to a measurably reduced antiviral, but not developmental function of ECT2. Fully resolving this issue requires actually understanding the precise molecular nature of both ECT2 functions. Nonetheless, we note that regardless of the exact interpretation of the reduced antiviral resistance in *te234* lines expressing ECT2^ΔN5^-mCherry, the result reinforces the conclusion reached from infections of *de25* and *vir-1* mutants that compromised antiviral resistance in *ect* mutants is not simply an indirect effect of developmental defects caused by dysfunction of the m^6^A-ECT axis.

**Figure 6.**
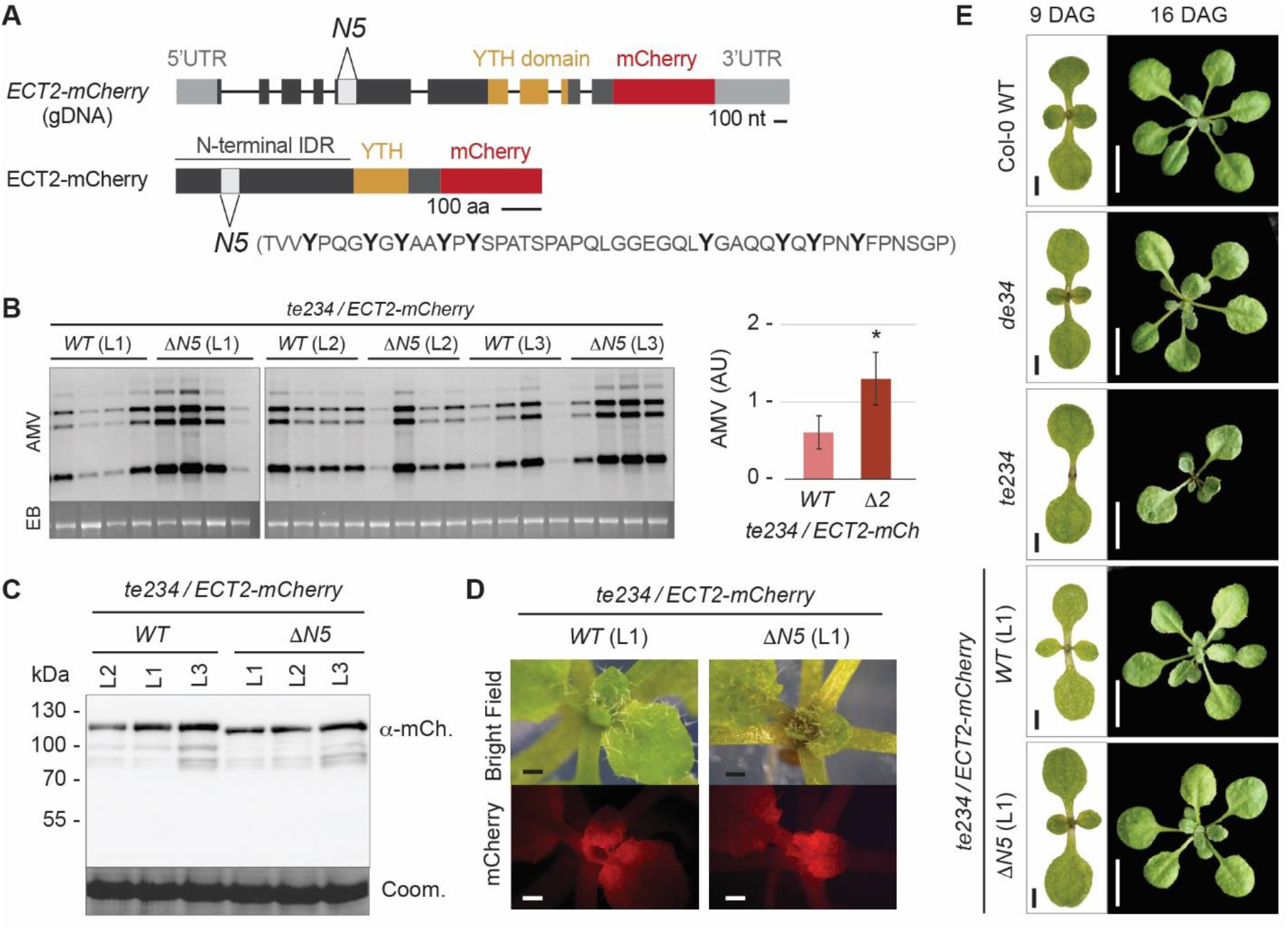
A Tyr-rich region of ECT2 IDR is necessary for antiviral but not for developmental functions of the protein. **(A)** Schematic representations of the annotated *ECT2* gene (AT3G13460.1, top) or protein (bottom) showing the deletion of the Tyr-rich region (ΔN5) in ECT2^ΔN5^-mCherry lines. **(B)** AMV systemic infection at 7 dpi of three independent complementation lines (L1-3) expressing ECT2-mCherry or ECT2^ΔN5^-mCherry in the *te234* background. Left, a representative RNA blot displaying AMV RNAs 1-4; right, the quantification histogram showing the mean of the three ECT2^WT^-mCherry (WT) or ECT2^ΔN5^-mCherry (ΔN5) lines. Ethidium bromide staining of rRNAs (EB) was used as RNA loading control. Error bars indicate standard deviations. AU, arbitrary units. Asterisk indicates significant differences from the WT (*: *p* < 0.05) applying Student’s t-test (n = 3). **(C)** Protein blot developed with mCherry antisera to show accumulation of ECT2^WT^-mCherry or ECT2^ΔN5^-mCherry fusion proteins in pools of the same samples used in panel B. Proteins on the membrane were stained with Coomassie-blue as loading control (lower panel). **(D)** Fluorescence microscopy of the second pair of true leaves of plants expressing ECT2^WT^-mCherry or ECT2^ΔN5^-mCherry in the *te234* background as indicated. Scale bars: 1 mm. **(E)** Phenotypes of seedlings and young rosettes of the indicated genotypes at 9 and 16 days after germination (DAG). Scale bars: 1 cm.

## DISCUSSION

### A new basal antiviral defense mechanism

Our analysis of the AMV-*Arabidopsis* interaction provides a clear case of the m^6^A-YTHDF axis acting as a basal antiviral defense layer: when the infection mechanism targeting the m^6^A-YTHDF axis is hampered by host *ALKBH9B* mutation, AMV systemic infection is severely disrupted and practically blocked in floral stems. However, systemic infectivity is restored upon additional mutation of *ECT2, ECT3*, and *ECT5*. These genetic data resemble the key arguments for the importance of RNAi as a basal antiviral defense mechanism in plants and insects (Deleris *et al*, 2006; Ding & Voinnet, 2007). Apart from obvious mechanistic questions that we touch on below, this discovery raises the important question of how widespread the use of this defense mechanism may be. AMV may be a special case, as it is one of only very few studied plant RNA viruses for which no anti-RNAi effector has been identified, and indeed prunus necrotic ringspot virus (PNRSV), a virus genetically and functionally closely related to AMV (Pallas *et al*, 2013), does not induce easily detectable siRNAs, unlike nearly all other studied plant RNA viruses (Herranz *et al*, 2015). On the other hand, a study comparing the *Arabidopsis* transcriptome in response to 11 plant viruses pointed out that differential expression of *ECT3* is a common response to all viruses, while the genes encoding the methylase components MTA and HAKAI1 and the YTHDF proteins ECT5 and ECT6 were differentially expressed upon infection with at least one virus (Postnikova & Nemchinov, 2012). This suggests that the m^6^A pathway is more generally implicated in plant-virus interactions. Clearly, it will be of key importance to analyze the degree to which RNA of other plant viruses contain m^6^A, what the effect of inactivation of genes encoding the major YTHDF proteins is on their infectivity, and whether they too employ mechanisms to interfere with the m^6^A-YTHDF axis. We note in this regard that studies on mammalian viruses have shown widespread implication of the m^6^A-YTHDF axis in animal-virus interactions, albeit not with the rigorous genetics employed here to demonstrate function as a first layer of defense. For example, depletion of mammalian YTHDF proteins was found to enhance viral replication of ZIKV (Lichinchi *et al*, 2016) and HIV-1 (Tirumuru *et al*, 2016; Jurczyszak *et al*, 2020), and to increase stability and protein expression of hepatitis B virus (HBV) (Imam *et al*, 2018) and endogenous retroviral mRNAs (Chelmicki *et al*, 2021). The m^6^A-YTHDF system is not universally employed for antiviral defense in animals, however, as viruses also appear to manipulate it for their benefit. For example, mammalian YTHDF proteins stimulate HIV-1 Gag processing and viral release (Tirumuru *et al*, 2016), and enhance viral translation and replication via binding to m^6^A-sites located in the 3’UTR of the viral RNAs (Kennedy *et al*, 2016). The difficulty of generalizing the role of m^6^A in viral RNA in host-virus interactions is also well illustrated by the case of HBV in which m^6^A in the 5’-UTR of viral pre-genomic RNA enhances reverse transcription, while m^6^A sites located in the 3’-UTR promote RNA destabilization (Imam *et al*, 2018).

### Are all m^6^A effects on AMV infectivity mediated by YTHDF proteins?

The marked restoration of AMV infectivity of *alkbh9b* mutants upon inactivation of *ECT2, ECT3*, and *ECT5* strongly argues that YTHDF proteins are the major, if not the exclusive, effectors of m^6^A-mediated basal AMV defense. In general, at 7 dpi, knockout of *ECT2/3/5* in the *alkbh9b* background leads to intermediate viral loads compared to *alkbh9b* and *ect2/ect3/ect5* mutants, although the outcome shows some variability. However, the most plausible explanation for this is that, in a hypermethylation context – where ALKBH9B is not functional – residual contributions by some of the remaining ECT proteins become relevant. Nonetheless, YTHDF-independent effects of m^6^A cannot be formally excluded at this point. For example, m^6^A may act as a secondary structure switch caused by weaker m^6^A-U compared to A-U base pairs (Kierzek & Kierzek, 2003; Liu *et al*, 2015), a property that may either benefit or impede viral replication. An evaluation of this potentially important possibility will require mapping of m^6^A sites in AMV to single-nucleotide resolution in *alkbh9b* and wild type backgrounds, as well as *in vivo* secondary structure analysis now feasible via sequencing-adapted methods (Poulsen *et al*, 2015).

### Source of adenosine methylation of AMV RNA

The observation that the infectivity of AMV is not drastically enhanced in *vir-1* mutants, in contrast to *te234*, constitutes strong evidence that the hypersusceptibility of composite *ect* mutants is not simply an indirect consequence of developmental defects in plants with a disabled m^6^A-YTHDF pathway. This important conclusion is reinforced by the hypersusceptibility of *ect2/ect5* mutants and of *te234* mutants expressing the ΔN5 deletion of ECT2, as both of these backgrounds do not show developmental delay and aberrant leaf morphology. VIR is required for adenosine methylation of endogenous mRNA (Růžička *et al*, 2017), but if it were also the case for AMV RNA, a hypersusceptibility phenotype would be expected, thus arguing that methylation of AMV RNA is VIR-independent. Such a scenario would be consistent with the cytoplasmic replication cycle of AMV, but nuclear localization of MAC/MACOM proteins in uninfected plants. It does, therefore, raise the pertinent question what the identity of the AMV RNA methyl transferase is. We see two possible explanations. Either AMV RNA methylation is carried out by an unidentified cytoplasmic adenosine methyltransferase, or, perhaps more likely, certain MAC/MACOM subunits translocate to the cytoplasm during AMV infection, as has been reported in different infection contexts in mammalian cells (Hao *et al*, 2019; Srinivas *et al*, 2021).

### Distinction between cellular and viral m^6^A-containing RNA and the basis of ECT-mediated antiviral defense

At first glance, it is a surprising result that ECT2/ECT3 carry out functions in both antiviral defense and in stimulation of cellular proliferation, especially given that endogenous mRNA targets of ECT2/ECT3 are enriched in factors involved in central metabolic processes such as oxidative phosphorylation and translation (Arribas-Hernández *et al*, 2021b). Thus, ECT2/ECT3 binding to endogenous targets presumably increases their expression, consistent with the predominant downregulation of ECT2/ECT3 mRNA targets in root meristem cells devoid of ECT2/ECT3/ECT4 activity (Arribas-Hernández *et al*, 2021b). By contrast, AMV RNA is repressed by ECT2/ECT3/(ECT4)/ECT5. What could dictate the different outcomes of binding of these proteins to endogenous mRNA versus AMV RNAs? As a working model to explain our results, we propose that m^6^A site multiplicity in AMV RNA may be a key factor distinguishing it from endogenous mRNA. Closely spaced m^6^A-sites in *cis* occupied by multiple ECT proteins would lead to high local concentrations of the IDRs, thus driving phase separation, and sequestration from the translation machinery. Such repression would not occur on endogenous mRNAs if they tend only to harbor single m^6^A sites. In this setting, the YTHDF proteins may use short linear motifs in their IDRs to recruit other cellular factors to enhance protein expression, but such activities would naturally be overridden upon condensation into a separate phase. Although admittedly speculative at this point, this model is consistent with a number of observations. First, abiotic stress indeed causes ECT2, ECT3 and ECT4 to condense into separate cytoplasmic bodies (Arribas-Hernández *et al*, 2018), identified as stress granules in the case of ECT2 in plants subjected to heat stress (Scutenaire *et al*, 2018), and purified ECT2 can separate into regularly shaped hydrogel-like particles *in vitro* (Arribas-Hernández *et al*, 2018). Thus, ECT2 has phase separation properties. Second, mammalian YTHDF2 has similar phase-separation properties *in vitro* that are markedly stimulated upon binding to RNA containing multiple, but not single, m^6^A sites (Ries *et al*, 2019; Fu & Zhuang, 2020; Gao *et al*, 2019). Third, phase separation depends on weak, polyvalent interactions between IDRs, often involving pi-pi, pi-charge, and hydrogen bonds, therefore singling out Tyr residues in IDRs as particularly strong drivers of phase separation (Vernon *et al*, 2018). It is noteworthy, therefore, that the short deletion in ECT2 that measurably impairs AMV defense, but not cellular proliferation, is strongly enriched in Tyr residues, and that such Tyr clusters are found in the IDRs of many plant YTHDF proteins, including ECT3 and ECT5 (**Supplemental Figure 6A, B**). Fourth, a recent computational approach to infer RNA modifications in plant-pathogenic viruses using high-throughput annotation of modified ribonucleotides (HAMR), a software that predicts modified ribonucleotides using high-throughput RNA sequencing data, revealed a higher proportion of RNA chemical modifications in comparison with mRNAs of Arabidopsis (Marquez-Molins *et al*, 2022). Finally, we note that although multiple m^6^A sites have been found in many endogenous mRNA targets across several species, there is as yet no evidence that these sites co-exist on the very same mRNA molecules. Indeed, in mammalian cells, the available evidence is to the contrary, as recent long-read analysis of mRNA from mammalian cells expressing a YTH domain fused to the C-to-U editing enzyme APOBEC showed preferential editing of only the C occurring in the methylated RR(m^6^A)CH context (Tegowski *et al*, 2022). We stress that despite the existence of evidence consistent with the proposed model for the basis of m^6^A-YTHDF-mediated basal AMV defense, we view this model as a conceptual framework of value in the design of future experiments to test its validity.

## ACKNOWLEDGMENTS

We thank Lorena Corachán, Freja Asmussen, and Lena Bjørn Johansson for their excellent technical assistance, and Joao Rato, Daniel Torrent-Silla, and the Bioinformatics Core Service at the Instituto de Biología Molecular y Celular de Plantas (IBMCP) for the support provided in the data analysis of the RNA differential expression analysis. Emilie Oksbjerg is thanked for running HyperTRIBE libraries in NextSeq sequencing system. And Kamil Růžička is thanked for seeds of *vir-1*. M.M.-P. was recipient of a Predoctoral Contract FPI-2015-072406 from the Subprograma FPI-MINECO (Formación de Personal Investigador– Ministerio de Economía y Competitividad). This work was supported by grant PID2020-115571RB-I00 to VP and FA from the Spanish MCIN/AEI/10.13039/501100011033 granting agency and Fondo Europeo de Desarrollo Regional (FEDER), by a project grant from Villum Fonden (Project #13397) and by an infrastructure grant from Carlsberg Fondet (CF18-1075) to PB. Part of this work was carried out in Prof. Brodersen’s lab at the University of Copenhagen (Denmark) thanks to a FEBS (Federation of European Biochemical Societies) Short-Term Fellowship to M.M.-P.

## AUTHOR CONTRIBUTIONS

M.M.-P. conceived the project, designed and performed the experiments, participated in the visualization of the results, and the data analysis and interpretation. F.A. participated in the experimental design and development, and the data analysis and interpretation. M.D.T. generated and characterized ECT2^ΔN5^-mCherry lines and contributed to generate ECT5 CRISPR mutants. S.R. performed bioinformatic analysis of HyperTRIBE data. L.A.-H. generated and characterized *te235*/*alkbh9b* lines, and participated in the visualization of the results, and the data analysis and interpretation. P.B and V.P. conceptualized the project and contributed to the experimental design, and the data analysis and interpretation. All authors corrected the manuscript and approved the final version.

## MATERIALS AND METHODS

### Plant growth conditions, virus inoculation, and northern blot analysis

The source and background of each Arabidopsis thaliana (Columbia-0) transgenic line are described in Supplemental Table 1. Plants were grown in 6 cm diameter pots in a growth chamber with a photoperiod of 25°C-16 h light/20°C-8 h dark. The mechanical inoculations of 15-19 days old plants were carried out using carborundum and purified virions (1 mg/mL) of AMV PV0196 isolate (Plant Virus Collection, DSMZ) in PE buffer (30 mM sodium phosphate buffer, pH 8). Next, detection of vRNAs was performed by northern blot analysis. Inoculated leaves and non-inoculated aerial tissue were harvested at the corresponding dpi. Tissues were grounded in liquid nitrogen with mortar and pestle and total RNA was extracted, following EXTRAzol reagent protocol (Blirt; Gdańsk, Poland), from 0.1 g leaf material. For northern blots, 500 ng of total RNA was denatured by formaldehyde treatment and after agarose gel electrophoresis in MOPS buffer, RNAs were transferred to positively charged nylon membranes (Roche; Basel, Switzerland) by capillarity in SSC buffer as previously described (Sambrook *et al*, 1988) and hybridized with digoxigenin-labeled riboprobes to detect AMV RNA 1, RNA 2, RNA 3 and sgRNA 4. Synthesis of the digoxigenin-labeled riboprobes, hybridization, and digoxigenin-detection procedures were performed as previously described (Pallás *et al*, 1998). Hybridization intensity signal was measured on files from Fujiflm LAS-3000 Imager using Fujiflm Image Gauge V4.0.

### Generation of Arabidopsis CRISPR/Cas9 transgenic lines

Arabidopsis lines carrying CRISPR/Cas9-mediated gene knockout were generated using the pKAMA-ITACHI Red (pKIR1.1) vector as it was previously described with some modifications (Tsutsui & Higashiyama, 2017). A simply adaptable single guide RNA (sgRNA) leads Cas9 endonucleases to target its complementary sequence, generating a DNA double-strand break (DSB) within a specific genomic sequence. The repair mechanisms are not completely efficient and often result in mutations (Lieber, 2010). In summary, different primers (**Supplemental Table 2**) were designed inside the ORF of ECT5 (AT3G13060), searching for protospacer adjacent motif (PAM) sequences (NGG), to work as sgRNAs. Then, ligations of the hybridized DNA oligomers with the vector, previously digested with AarI enzyme, were performed. Agrobacterium tumefaciens GV3101 was transformed with the resulting vectors, and cultures expressing two different sgRNAs were used to generate Arabidopsis stable transgenic lines by floral dip transformation (Clough & Bent, 1998) in *ect2-1* and *de23* backgrounds. Selection of T1 transgenic plants was carried out by hygromycin resistance and a first assortment of Cas9-free T2 seeds was performed by absence of red fluoresce. Final genotyping by PCR with specific primers (**Supplemental Table 2**) was realized from T2 plants and confirmed in T3 transformants. Finally, an RT-PCR with specific primers (**Supplemental Table 2**) of total RNA extraction from plants of selected lines and the sequencing of the cDNA products verified the non-frameshift mutations (**Supplemental Figure 3C, D**).

### Generation of Arabidopsis ECT2^ΔN5^-mCherry lines

Cloning and line selection of ECT2^ΔN5^-mCherry was performed by USER cloning (Bitinaite and Nichols, 2009) and agrobacterium-mediated transformation as in (Arribas-Hernández *et al*, 2018). In brief, USER-compatible primers (LA336, MT5, MT6, LA337) were used to amplify fragments from *ECT2Pro:ECT2gDNA-mCherry:ECT2Ter*; a construct previously generated by Arribas-Hernandez et al., 2018. The fragments were purified and inserted into pCAMBIA3300U (pCAMBIA3300 with a double PacI USER cassette inserted between the PstI-XmaI sites at the multiple cloning site) (Nour-Eldin *et al*, 2006) after which the plasmid was transformed into Escherichia coli DH5α (NEB). Kanamycin resistant colonies were selected and their plasmids purified followed by restriction digestion and sequencing prior to introduction into Agrobacterium tumefaciens GV3101 for plant transformation. Arabidopsis stable transgenic lines were generated by floral dip transformation (Clough & Bent, 1998) of *ect2-1/ect3-1/ect4-2* (*te234*) (Arribas-Hernández *et al*, 2018), and selection of primary transformants (T1) was done on MS-agar plates supplemented with glufosinate-ammonium (Merck, Darmstadt, Alemania) (10 mg/L). T2 generation was used for the infection experiments.

### RNA differential expression analysis between Arabidopsis Col-0 WT MOCK and AMV samples

2-weeks Arabidopsis Col-0 WT plants were mechanically inoculated with PE buffer (MOCK plants) or AMV PV0196 isolate viral particles in PE. Tissues from the young, emerging rosette leaves were harvested at 8 dpi and grounded in liquid nitrogen with mortar and pestle. Total RNA was extracted from 0.1 g leaf material using EXTRAzol reagent protocol (Blirt; Gdańsk, Poland) and treated with DNase I for 30 min at 37ºC. The experiment consisted of three biological replicates (8 individual plants/ replicate) of MOCK and AMV WT inoculated plants. Generation and sequencing of the cDNA libraries were performed by the Genomic Service (SCSIE) of the Universidad de Valencia. Six TruSeq Stranded cDNA libraries (three for healthy and three for AMV-infected plants) were sequenced by non-paired end sequencing (75 bp) in a NextSeq 550 (Illumina; San Diego, California, USA). The bioinformatics analysis was carried out by the Bioinformatic Service of IBMCP. DESeq2 package was the tool used for the differential expression analysis and a threshold of 1RPMK was applied.

### Quantitative reverse-transcription PCR (qPCR)

Four μl of the reaction containing DNA-free RNA were mixed with 0.5 μg (1 μl) of oligo(dT)18 (ThermoFisher Scientific; Waltham, Massachusetts, USA), denatured for 5 min at 65ºC and added to a total of 20 μl of first-strand cDNA synthesis reaction containing 1 U of RevertAid H Minus Reverse Transcriptase, 0.5 U of Ribolock, 4 mM dNTP mix and 1x RT buffer (ThermoFisher Scientific; Waltham, Massachusetts, USA). The RT reaction proceeded at 42°C for 1 hour, followed by 10 min at 70°C to deactivate the enzyme. qPCR reactions were carried out using PyroTaq EvaGreen mix Plus (ROX; CulteK Molecular Bioline, Madrid, Spain) according to the manufacturer’s instructions. All analyses were performed in triplicate on an ABI 7500 Fast-Real Time qPCR instrument (Applied Biosystems, Waltham, MA, USA). Specific oligonucleotides efficiencies were tested by qPCR using 10-fold serial dilutions of the corresponding cDNA (**Supplemental Table 2**). Expression analysis was performed by calculating the relative quantification (RQ) values with respect to the endogenous genes *F-BOX* and *PDF2* (Lilly *et al*, 2011) of three biological replicates. Student’s t-test was applied to ΔCt values for statistical analysis.

### Fluorescence microscopy

Imaging of plants at the rosette was done using a Leica MZ16 F fluorescence stereomicroscope, as described by Arribas-Hernández et al. (2018).

### Analysis of ECT2-RNA^AMV^ interaction by HyperTRIBE

Five independent lines expressing ECT2 fused to the catalytic domain of the adenosine deaminase acting on RNA (ADAR) on the *ect2-1* background and five control lines expressing only ADAR on WT background (Arribas-Hernández *et al*, 2021a) were inoculated at 16 days with AMV (McMahon *et al*, 2016; Xu *et al*, 2018). Tissue from the innermost, young rosette leaves of each line (10 plants per sample) was selected at 7 dpi and grounded in liquid nitrogen with mortar and pestle. After total RNA extraction, precipitation of nucleic acids was carried out with 3,75 M LiCl (−20ºC, overnight) to avoid tRNA precipitation. Next, these samples were submitted to a ribosomal RNA depletion treatment and cDNA libraries were prepared with CORALL Total RNA-Seq Library Prep Kit. The libraries were sequenced by paired end sequencing (150 pb) on an Illumina NEXTseq 550 platform with v2.5 flow cell (300 cycles). Libraries were first quality checked before being merged across lanes, then trimmed, duplicated, and mapped using the CORALL Total RNA-Seq integrated Data Analysis Pipeline. Quantifications of reads mapping to AMV and ADAR were done using Salmon (Patro *et al*, 2017) using paired-end stranded mode. Mapped .bam files were converted to reflect forward and reverse strands before processing using hyperTRIBER (Rennie *et al*, 2021): briefly, base counts were aggregated from reads for all positions containing at least one mismatch, which were subsequently filtered for viral positions containing putative A-G edits in at least two of five lines expressing ECT2-FLAG-ADAR. 395 candidates were formally tested for differential occurrence of the base G while adjusting for differences in both read coverage at the position for each condition and variation in the levels of the ECT2-FLAG-ADAR. The results were subsequently FDR corrected in R, and the R function cor.test was used to calculate Pearson’s correlation between ADAR and editing proportions (A/(A+G)) in ECT2-FLAG-ADAR samples. All plots were produced in R (R Core Team, 2021).

### Western blot analysis

Total protein was extracted homogenizing 0.1 g of tissue from 7 dpi infected plants (same samples used for northern blot analysis in Figure 6B) with 3 volumes of extraction buffer (0.4 M Tris-HCl pH 8.8, 2% SDS, 15% glycerol, 0.1 M DTT), and heat at 95ºC for 10 min. After centrifugation, 5 µL of Laemmli 6x (0.3 M Tris-HCl pH 6.8, 10% SDS, 0.05% xylene cyanol/ bromophenol blue, 15% β-mercaptoethanol) were added to 25 µL of each sample and loaded into a 10% SDS-PAGE gel. The electrophoresis was run at 100 V for around 2 h, and proteins were transferred to a PVDF membrane, previously activated with methanol, at 30 V and 4ºC overnight. ECT2-mCherry protein was detected using the anti-mCherry antibody (ab183628, abcam; Cambridge, United Kindom) at dilution 1:10000. Secondary antibody and detection procedure was carried out following the manufacturer’s instructions (ECL™ Prime Western Blotting System, Merck; Darmstadt, Germany).

## DATA AVAILABILITY

RNA-Seq and HyperTRRIBE RNA-Seq data: PRJEB56577 (https://www.ebi.ac.uk/ena/browser/view/PRJEB56577).

## SUPPLEMENTAL INFORMATION TITLES AND LEGENDS

**Supplemental Figure 1. Single mutation of *ECT2, 3, 4* or *5* alone does not alter AMV infection. (A)** *ECT5* gene model (AT3G13060.2) showing the location of the *ect5-1* T-DNA insertion (SALK_131549, **Supplemental Table 1**). Exons are depicted as boxes and introns as lines. Untranslated regions (UTRs) are colored in light grey, and the exonic regions encoding the YTH domain are highlighted in light brown. Position of the primers used to amplify full length *ECT5* are depicted in green, and genotyping primers in blue. **(B)** Ethidium bromide-stained agarose gels showing *ect5-1* genotyping by PCR (left panel) to corroborate the presence of the T-DNA, and RT-PCR (right panel) to amplify the full-length transcript. Location of primers LP (left primer, genomic), RP (right primer, genomic), and LB (left border primer, T-DNA) are schematically shown in blue (A), whereas nucleotide sequences can be found in **Supplemental Table 2. (C)** RNA blot analysis of AMV systemic infection at 7 dpi in *ect2-1, ect3-1, ect4-2* (left panels), and *ect5-1* (right panels) mutants compared to wild type plants. Each panel shows a representative RNA blot displaying AMV RNAs 1-4 (left) and its quantification histogram (right). Ethidium bromide staining of rRNAs (EB) was used as RNA loading control. Error bars indicate standard deviation, and asterisks indicate *p* < 0.05 relative to wild type and applying Student’s t-test (n = 4). AU, arbitrary units.

**Supplemental Figure 2. Susceptibility of *ect2/ect3, ect2/ect3/ect4* and *ect2/ect3/ect5* mutants to AMV infection (extended data) (A)** RNA blot analysis of AMV systemic infection at 6 dpi in mutants with two allele combinations of *ect2/ect3, de23* (*ect2-1/ect3-1*) and *Gde23* (*ect2-3/ect3-2*), compared to wild type plants. Left, blot displaying AMV RNAs 1-4; right, quantification histogram. Dashed lines indicate non-contiguous samples that are analyzed on the same membrane. Ethidium bromide staining of rRNAs (EB) was used as RNA loading control. **(B, C)** Quantification of three independent experiments comparing AMV systemic infection levels in *de23* and *te234* (B) or *te235* (C) by RNA blot analysis. The height of the bars represents the average, for the 3 experiments, of the means over four samples in each experiment. In all cases, error bars indicate standard deviation, and asterisks (**) show *p* < 0.01 relative to wild type applying Student’s t-test. AU, arbitrary units.

**Supplemental Figure 3. *ect5* CRISPR/Cas9 mutants on the *ect2-1* and *de23* backgrounds. (A)** Schematic map of the five nuclear chromosomes (Chr 1-5) of *Arabidopsis thaliana*. The position of the thirteen genes encoding YTH domain proteins (eleven YTHDFs (ECTs) and two YTHDCs (CPSF30 and AT4G11970)) is indicated. A black arrow marks the positions of *ECT2* and *ECT5* loci, in close proximity. Mb, megabase pairs. **(B)** Schematic representations of the annotated *ECT5* gene (AT3G13060.2) showing the location of the *ect5-1* T-DNA insertion (SALK_131549) and, marked in blue, the four deletions engineered using CRISPR/Cas9 in the *ect2-1* (*ect5-2* and *ect5-3*) or *de23* (*ect5-4* and *ect5-5*) backgrounds. Exons are depicted as boxes and introns as lines. Untranslated regions (UTRs) are colored in light gray, and the exonic regions encoding the YTH domain are highlighted in light brown. **(C)** Alignment between the Sanger-sequencing reads obtained from cDNA of the four CRISPR/Cas9 alleles described in B, and wild type *ECT5* cDNA (AT3G13060.2) showing the deletions produced by Cas9. **(D)** Alignment between the amino acid sequences of wild type ECT5 and the proteins predicted to be encoded by the Cas9-edited *ect5* loci described in B and C. The four deletions cause shifts in the ORF that result in short amino acid stretches (marked in red) that end with a premature stop codon (*).

**Supplemental Figure 4. AMV susceptibility in independent *ect2/ect5* and *ect2/ect3/ect5* transgenic lines**. RNA blot of AMV systemic infection at 6 dpi in *de23, ect2-1/ect5-3* and *ect2-1/ect3-1/ect5-5* lines compared to wild type. Left, RNA blot displaying AMV RNAs 1-4; right, quantification histogram. Error bars indicate standard deviation, and asterisks (**) indicate *p* < 0.01 relative to wild type applying Student’s t-test. Ethidium bromide staining of rRNAs (EB) was used as RNA loading control. AU, arbitrary units.

**Supplemental Figure 5. Editing proportions (G/(A+G)) for the top three most significant positions in AMV RNAs in individual samples**. Histograms showing the editing proportions (A-to-G) of the three top positions in AMV RNAs identified by HyperTRIBE pipeline over five independent transgenic lines of ECT2-FLAG-ADAR and FLAG-ADAR control lines.

**Supplemental Figure 6. Analysis of ECT2**^**ΔN5**^**-mCherry mutants (extended data). (A)** Tyr-rich region deleted in ECT2^ΔN5^-mCh mutants is conserved to some extent in ECT3, 4 and 5 (shaded in blue). Multiple sequence alignment of the N-terminal 200 amino acid region of these proteins (Clustal Omega). **(B)** Amino acids proportion (%) of ECT2, 3, 4 and 5 in the section that aligned with the Tyr-rich region **(C)** Full complementation of the developmental phenotype of *te234* mutants by expression of ECT2^ΔN5^-mCherr*y* is comparable to that of wild type ECT2^WT^-mCherry, as seen three independent lines of each type (used for the infection assays in **Figure 6B**) compared to wild type or *de34* plants. DAG, days after germination. Scale bars are 1mm for the upper panels, and 1 cm for the lower.

