## Supplemental Figures and tables for "Plant YTHDF proteins are direct effectors of antiviral immunity against an m^6^A-containing RNA virus"

**Supplemental Table 1. List of *Arabidopsis thaliana* mutant lines employed in this work.** All the lines are in the Col-0 ecotype.

| ID | Genotype | Methodology | Ref * |
|---|---|---|---|
| **Single KO mutants** | | | |
| ***ect2-1*** | SALK_002225 | T-DNA insertion | 1 |
| ***ect3-1*** | SALK_077502 | T-DNA insertion | 1 |
| ***ect4-2*** | GK_241H02 | T-DNA insertion | 1 |
| ***ect5-1*** | SALK_131549 | T-DNA insertion | this work |
| ***ect2-3*** | GK_132F02 | T-DNA insertion | 1 |
| ***ect3-2*** | GABIseq_487H12 | T-DNA insertion | 1 |
| **Double and triple KO mutants** | | | |
| ***de23*** | *ect2-1/ect3-1* | Genetic cross | 1 |
| ***Gde23*** | *ect2-3/ect3-2* | Genetic cross | 1 |
| ***de24*** | *ect2-1/ect4-2* | Genetic cross | 1 |
| ***de34*** | *ect3-1/ect4-2* | Genetic cross | 1 |
| ***te234*** | *ect2-1/ect3-1/ect4-2* | Genetic cross | 1 |
| ***de25*** | *ect2-1/ect5-2* | CRISPR/Cas9-deletion of *ECT5* in *ect2-1* | this work |
| ***ect2-1/ect5-3*** | *ect2-1/ect5-3* | CRISPR/Cas9-deletion of *ECT5* in *ect2-1* | this work |
| ***te235*** | *ect2-1/ect3-1/ect5-4* | CRISPR/Cas9-deletion of *ECT5* in *de23* | this work |
| ***ect2-1/ect3-1/ect5-5*** | *ect2-1/ect3-1/ect5-5* | CRISPR/Cas9-deletion of *ECT5* in *de23* | this work |
| ***alkbh9b/te235*** | *alkbh9b/ect2-1/ect3-1/ect5-4* | Genetic cross | this work |
| **Transgenic lines** | | | |
| ***de23/ECT2-mCh*** **(2 lines)** | *ect2-1/ect3-1* *ECT2p:ECT2-mCherry-ECT2t* | USER cloning & Agrob. transformation | 1 |
| ***de23/ECT2$^{W464A}$-mCh*** **(2 lines)** | *ect2-1/ect3-1* *ECT2p:ECT2$^{W464A}$-mCherry-ECT2t* | USER cloning & Agrob. transformation | 1 |
| ***ect2-1*/ECT2-FLAG-ADAR (5 lines)** | *ect2-1* *ECT2p:ECT2-FLAG-ADAR-ECT2t* | USER cloning & Agrob. transformation | 2 |
| **FLAG-ADAR (5 lines)** | *ECT2p: FLAG-ADAR-ECT2t* | USER cloning & Agrob. transformation | 2 |
| ***te234/ECT2-mCh*** **(3 lines)** | *ect2-1/ect3-1/ect4-2* *ECT2p:ECT2-mCherry-ECT2t* | USER cloning & Agrob. transformation | 1 |
| ***te234/ECT2$^{\Delta2}$-mCh*** **(3 lines)** | *ect2-1/ect3-1/ect4-2* *ECT2p:ECT2$^{\Delta2}$-mCherry-ECT2t* | USER cloning & Agrob. transformation | this work |

* References: Arribas-Hernández et al., 2018[1]; Arribas-Hernández et al., 2021[2]

**Supplemental Table 2.** List of oligonucleotides used in this work.

| | | |
|---|---|---|
| qPCR | ECT5_qPCR_F | AGGCAGCAAAGAAGCAGTCA |
| | ECT5_qPCR_R | CCAGACCCGGTCAGACGTAA |
| | MTA_qPCR_F | CGGATCCACCATGGGACATT |
| | MTA_qPCR_R | AAGCTCCAAACATTCACGGC |
| | MTB_qPCR_F | GTACTGTGTTTCAGCGTTCC |
| | MTB_qPCR_R | TTCTGAGTCGAACCATAAGGAG |
| | VIR_qPCR_F | ACGCAAGTCCAGCCTTACTATCAC |
| | VIR_qPCR_R | CGGTCACTTAATAGAGCCTGAATGG |
| *ect5* genotyping | LPect5-1 | TGGACACGGTTCTTTCTCATC |
| | RPect5-1 | TGATCAAATCAGGTTGGCTTC |
| | LB | ATTTTGCCGATTTCGGAAC |
| | ECT5_F | GGTCTCcCATGGCAACGACTCAATCACAC |
| | ECT5_R | GGTCTCgCTAGTTGGATTGACCAGAAGAG |
| CRISPR/Cas9 RNA guides | sgRNA_ECT5.1F | ATTGAAGTGTCCGGCAAAATCAC |
| | sgRNA_ECT5.1R | AAACGTGATTTTGCCGGACACTT |
| | sgRNA_ECT5.2F | ATTGTGAAAACCCCACAGGCGAA |
| | sgRNA_ECT5.2R | AAACTTCGCCTGTGGGGTTTTCA |
| | sgRNA_ECT5.3F | ATTGGGCAAGGGACTTGCTGCAG |
| | sgRNA_ECT5.3R | AAACCTGCAGCAAGTCCCTTGCC |
| CRISPR/Cas9 genotyping | C9_ECT5g_F | GCAGACTGTCCCGGCAAAC |
| | C9_ECT5g_R | CGGGCTGAGTTGGAGAAGTAA |
| | C9(LA)_ECT5-F | CTATGTTGGGGAGCCAGG |
| | C9(LA)_ECT5-R | GCTGGTAATACGGTGATGC |

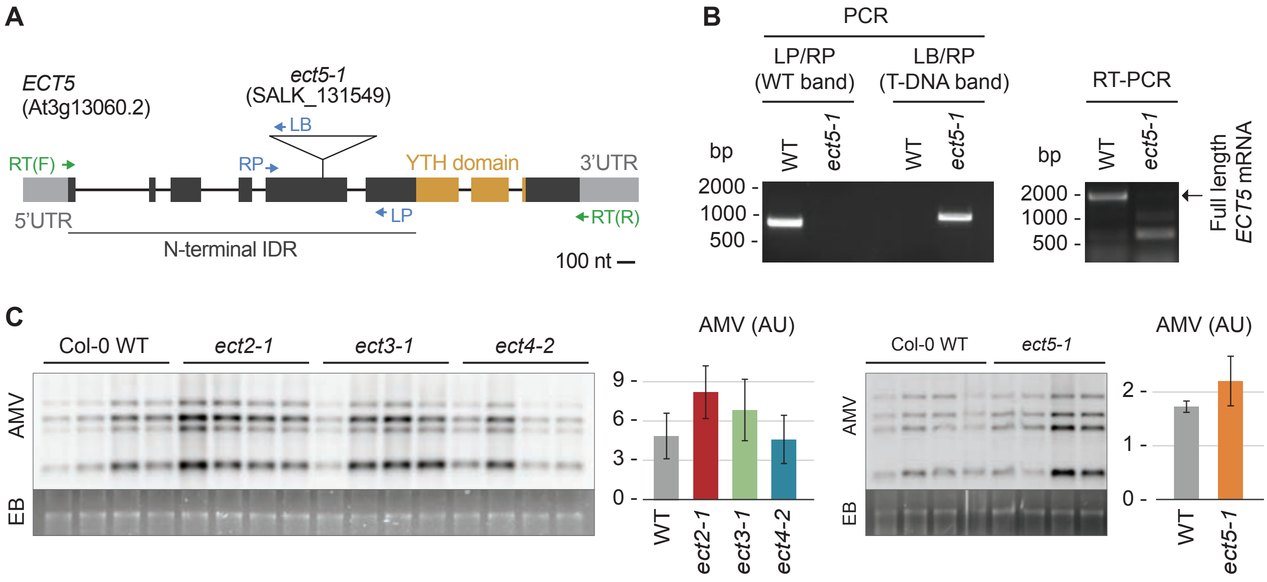


**Supplemental Figure 1. Single mutation of *ECT2*, *3*, *4* or *5* alone does not alter AMV infection.**
**(A)** *ECT5* gene model (AT3G13060.2) showing the location of the *ect5-1* T-DNA insertion (SALK_131549, **Supplemental Table 1**). Exons are depicted as boxes and introns as lines. Untranslated regions (UTRs) are colored in light grey, and the exonic regions encoding the YTH domain are highlighted in light brown. Position of the primers used to amplify full length *ECT5* are depicted in green, and genotyping primers in blue. **(B)** Ethidium bromide-stained agarose gels showing *ect5-1* genotyping by PCR (left panel) to corroborate the presence of the T-DNA, and RT-PCR (right panel) to amplify the full-length transcript. Location of primers LP (left primer, genomic), RP (right primer, genomic), and LB (left border primer, T-DNA) are schematically shown in blue in panel A, whereas nucleotide sequences can be found in **Supplemental Table 2**. **(C)** RNA blot analysis of AMV systemic infection at 7 dpi in *ect2-1*, *ect3-1*, *ect4-2* (left panels), and *ect5-1* (right panels) mutants compared to wild type plants. Each panel shows a representative RNA blot displaying AMV RNAs 1-4 (left) and its quantification histogram (right). Ethidium bromide staining of rRNAs (EB) was used as RNA loading control. Error bars indicate standard deviation, and asterisks indicate $p < 0.05$ relative to wild type and applying Student's t-test (n = 4). AU, arbitrary units.

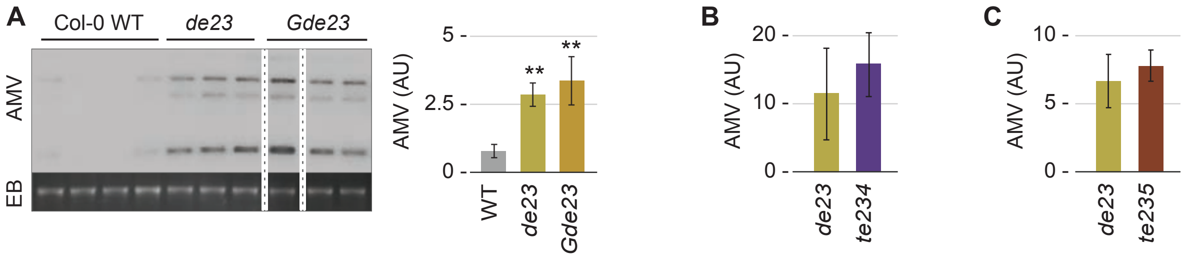


**Supplemental Figure 2. Susceptibility of *ect2/ect3*, *ect2/ect3/ect4* and *ect2/ect3/ect5* mutants to AMV infection (extended data). (A)** RNA blot analysis of AMV systemic infection at 6 dpi in mutants with two allele combinations of *ect2/ect3*, *de23* (*ect2-1/ect3-1*) and *Gde23* (*ect2-3/ect3-2*), compared to wild type plants. Left, blot displaying AMV RNAs 1-4; right, quantification histogram. Dashed lines indicate non-contiguous samples that are analyzed on the same membrane. Ethidium bromide staining of rRNAs (EB) was used as RNA loading control. **(B, C)** Quantification of three independent experiments comparing AMV systemic infection levels in *de23* and *te234* (B) or *te235* (C) by RNA blot analysis. The height of the bars represents the average, for the 3 experiments, of the means over four samples in each experiment. In all cases, error bars indicate standard deviation, and asterisks (**) show $p < 0.01$ relative to wild type applying Student's t-test. AU, arbitrary units.

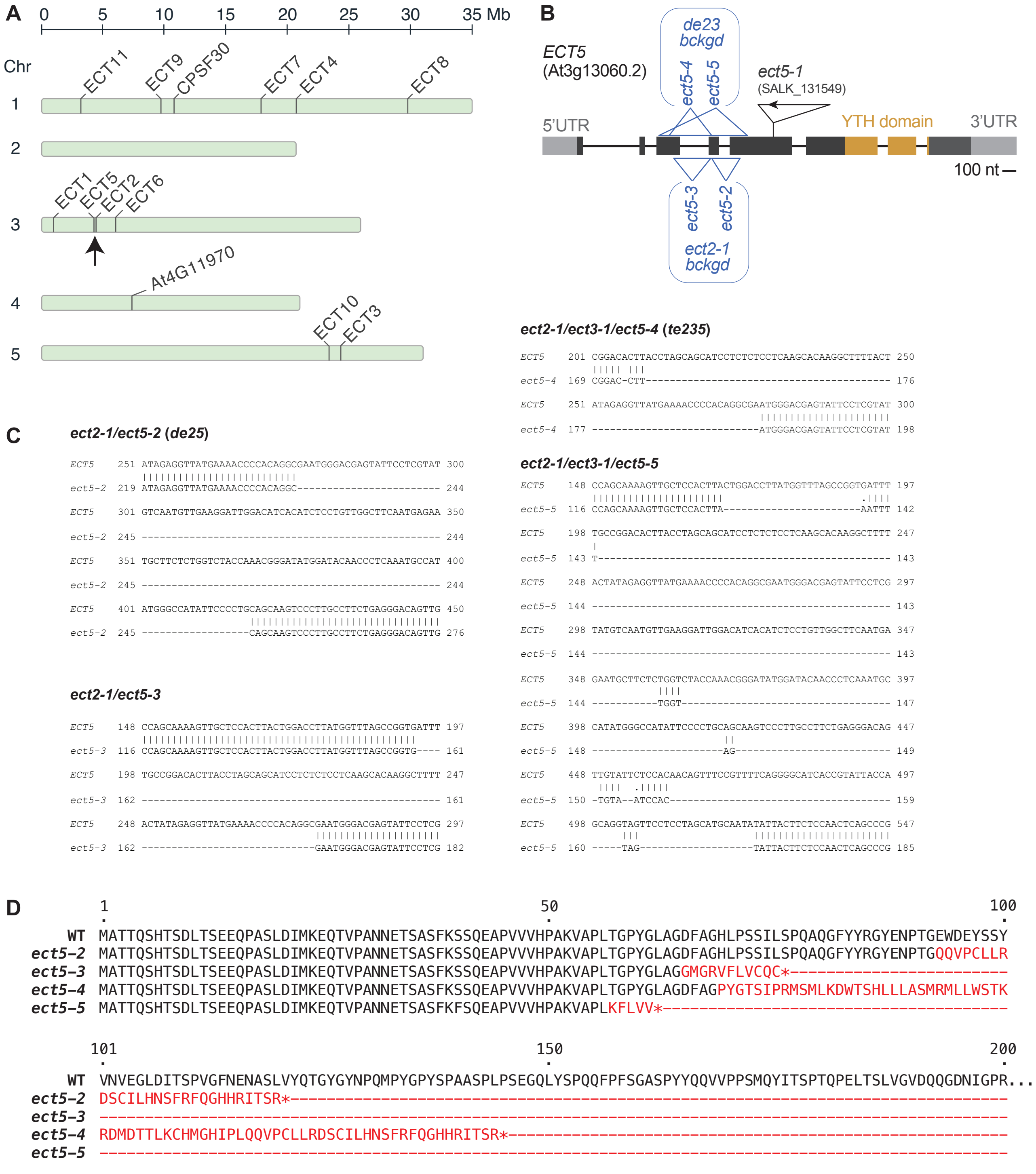


**Supplemental Figure 3.** ***ect5*** **CRISPR/Cas9 mutants on the** ***ect2-1*** **and** ***de23*** **backgrounds.**
**(A)** Schematic map of the five nuclear chromosomes (Chr 1-5) of *Arabidopsis thaliana*. The position of the thirteen genes encoding YTH domain proteins (eleven YTHDFs (ECTs) and two YTHDCs (CPSF30 and AT4G11970)) is indicated. A black arrow marks the positions of *ECT2* and *ECT5* loci, in close proximity. Mb, megabase pairs. **(B)** Schematic representations of the annotated *ECT5* gene (AT3G13060.2) showing the location of the *ect5-1* T-DNA insertion (SALK_131549) and, marked in blue, the four deletions engineered using CRISPR/Cas9 in the *ect2-1* (*ect5-2* and *ect5-3*) or *de23* (*ect5-4* and *ect5-5*) backgrounds. Exons are depicted as boxes and introns as lines. Untranslated regions (UTRs) are colored in light gray, and the exonic regions encoding the YTH domain are highlighted in light brown. **(C)** Alignment between the Sanger-sequencing reads obtained from cDNA of the four CRISPR/Cas9 alleles described in B, and wild type *ECT5* cDNA (AT3G13060.2) showing the deletions produced by Cas9. **(D)** Alignment between the amino acid sequences of wild type *ECT5* and the proteins predicted to be encoded by the Cas9-edited *ect5* loci described in B and C. The four deletions cause shifts in the ORF that result in short amino acid stretches (marked in red) that end with a premature stop codon (*).

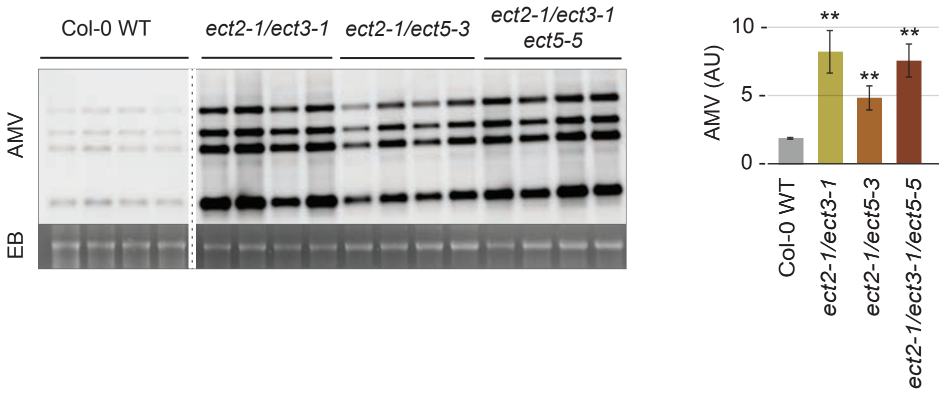


**Supplemental Figure 4. AMV susceptibility in independent *ect2/ect5* and *ect2/ect3/ect5* transgenic lines.** RNA blot of AMV systemic infection at 6 dpi in *de23*, *ect2-1/ect5-3* and *ect2-1/ect3-1/ect5-5* lines compared to wild type. Left, RNA blot displaying AMV RNAs 1-4; right, quantification histogram. Error bars indicate standard deviation, and asterisks (**) indicate $p < 0.01$ relative to wild type applying Student's t-test. Ethidium bromide staining of rRNAs (EB) was used as RNA loading control. AU, arbitrary units.

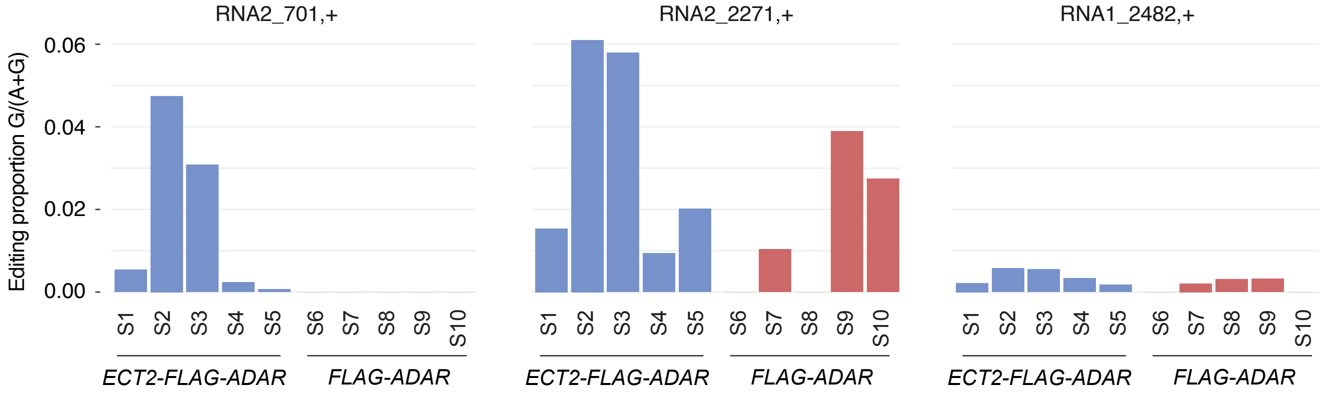


**Supplemental Figure 5. Editing proportions (G/(A+G)) for the top three most signi¬ficant positions in AMV RNAs in individual samples.** Histograms showing the editing proportions (A-to-G) of the three top positions in AMV RNAs identified by HyperTRIBE pipeline over five independent transgenic lines of ECT2-FLAG-ADAR and FLAG-ADAR control lines.

**A**

```
ECT3      ----MANPDHVSDVLHNLSIDPTTKALAPDSETKLQGAYGGNGNDFLLNDELVEATKIGK      56
ECT5      MATTQSHTSDLTSEEQPASLDIM-KEQTVPANNETSASFKSSQEAPVVVHPAKVAPLTGP      59
ECT2      MATVAPPADQATDLLQKLSLDSPAKASEIPEPNKKTAVYQYGG---VDVH-GQV------      50
ECT4      MSTVAPPADQAADVLKKLSLDSKSRTLEIPEPTKKTGVYQYGA---MDSN-GQV------      50
                  .. :.  :  *:*   :       .:  . :  .    :  .

ECT3      PSLLSKDGGVTKDKGSNLKKLGYQSAAYNAKGSYGKGAYAYGYYPPAYQYPRHGYTGSYA      116
ECT5      YGLA---GDFAGHLPSSILSP------------QAQGFYYRGYENPTGEW----------      94
ECT2      PSYD---RSLTPMLPSDAADP---SVCYV-P-NPYNPYQYYNVYGSGQEW----------      92
ECT4      PSFD---RSLSPMLPSDALDP---SVFYV-P-NVYQQPYYYG---YG-------------      86
           .      ..:    *.  .               :     .

ECT3      SGKTNLQY-QYLTQQGRSAGNGQSYGGYMDNIYSNYGMCGPYTNGYGYGS-YGYDSWK--      172
ECT5      -----DEYSSYVNVEGLDITSPVGFNENA---------SLVYQTGYGYNPQMPYGPYSPA      140
ECT2      -----TDYPAYTNPEGVDMNSG-IYGENG---------TVVYPQGYGYAA-YPYSPATSP      136
ECT4      -----SDYTGYTNSESVDMTSG-AYGENA---------SLVYPQGYGYAA-FPYSPATSP      130
                :*  * . :. .  .   :.               *  ****     *.  .

ECT3      ----------YMPNWYAVNNT----------Y--------KPRNG------YHGYG----      194
ECT5      ASPLPSEGQLYSPQQFPFSGASPYYQQVVPPSMQYITSPTQPELTSL-------VGVDQQ      193
ECT2      APQLGGEGQLYGAQQYQYPNYF----PNSGPYASSVATPTQPDLSANKPAGVKTL--PAD      190
ECT4      APQLGGDGQLYGAQQYQYPFPL---TASSGPFASSVPASTQSKLSTNKAANSASAGIPKG      187
                    *  : :                        :

ECT3      KENIEG-------       200
ECT5      GDNIGPR------       200
ECT2      SNNVASAAGI---       200
ECT4      MNGSAPVKPLNQS       200
           :.
```

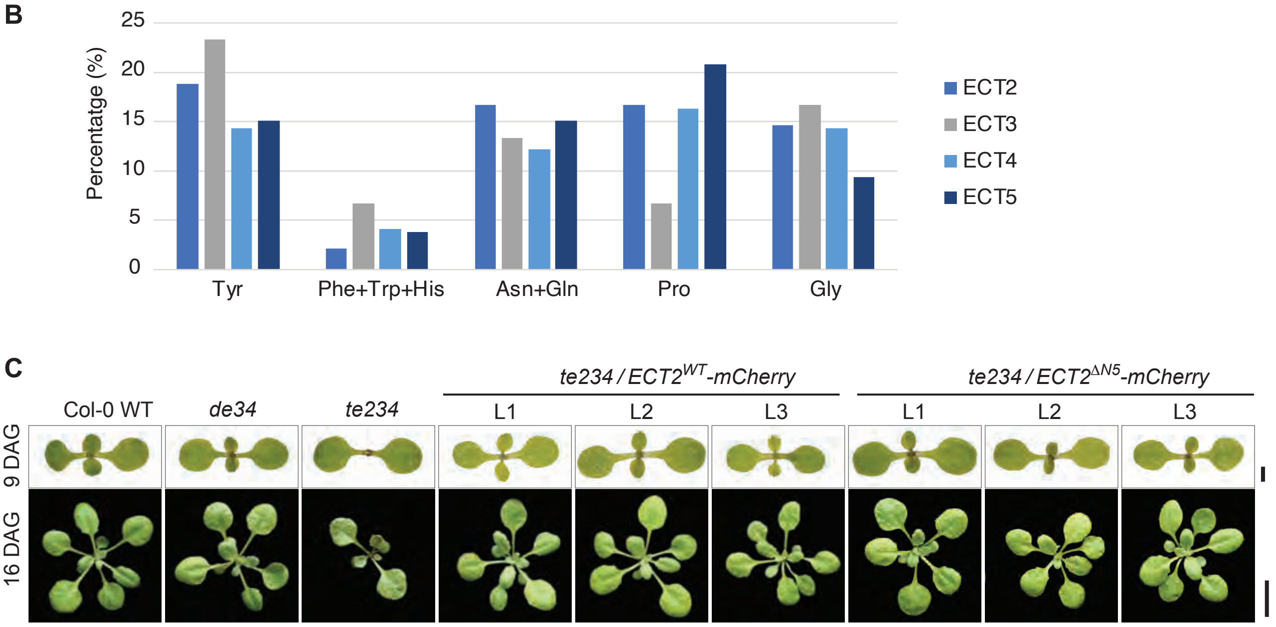


**Supplemental Figure 6. Analysis of ECT2$^{\Delta N5}$-mCherry mutants (extended data). (A)** Tyr-rich region deleted in ECT2$^{\Delta N5}$-mCh mutants is conserved to some extent in ECT3, 4 and 5 (shaded in blue). Multiple sequence alignment of the N-terminal 200 amino acid region of these proteins (Clustal Omega). **(B)** Amino acids proportion (%) of ECT2, 3, 4 and 5 in the section that aligned with the Tyr-rich region **(C)** Full complementation of the developmental phenotype of *te234* mutants by expression of ECT2$^{\Delta N5}$-mCherry is comparable to that of wild type ECT2$^{WT}$-mCherry, as seen three independent lines of each type (used for the infection assays in **Figure 6B**) compared to wild type or *de34* plants. DAG, days after germination. Scale bars are 1 mm for the upper panels, and 1 cm for the lower.
